# Circadian Proteins Cry and Rev-erb Converge to Deepen Cellular Quiescence by Downregulating Cyclin D and Cdk4,6

**DOI:** 10.1101/2021.07.30.454549

**Authors:** Xia Wang, Bi Liu, Qiong Pan, Jungeun Sarah Kwon, Matthew A. Miller, Kimiko Della Croce, Guang Yao

## Abstract

The proper balance and transition between cellular quiescence and proliferation are critical to tissue homeostasis, and their deregulations are commonly found in many human diseases, including cancer and aging. Recent studies showed that the reentry of quiescent cells to the cell cycle is subjected to circadian regulation. However, the underlying mechanisms are largely unknown. Here, we report that two circadian proteins, Cryptochrome (Cry) and Rev-erb, deepen cellular quiescence in rat embryonic fibroblasts, resulting in stronger serum stimulation required for cells to exit quiescence and reenter the cell cycle. This finding was opposite from what we expected from the literature. By modeling a library of possible regulatory topologies linking Cry and Rev-erb to a bistable Rb-E2f gene network switch that controls the quiescence-to-proliferation transition and by experimentally testing model predictions, we found Cry and Rev-erb converge to downregulate Cyclin D/Cdk4,6 activity, leading to an ultrasensitive increase of the serum threshold to activate the Rb-E2f bistable switch. Our findings suggest a mechanistic role of circadian proteins in modulating the depth of cellular quiescence, which may have implications in the varying potentials of tissue repair and regeneration at different times of the day.

## INTRODUCTION

Upon growth signals, various types of quiescent cells (e.g., adult stem and progenitor cells) can reenter the cell cycle to proliferate. This quiescence-to-proliferation transition is fundamental to tissue homeostasis and repair ^1–3^. This transition, as shown in recent studies, also appears to be affected by the circadian clock. For example, quiescent neural stem cells and progenitor cells initiate neurogenesis to produce new neurons in a circadian-dependent manner ^4, 5^; similarly does the circadian activation of hair follicle stem cells for tissue renewal ^6^.

Circadian clocks are present in cells throughout mammalian tissues. These clocks are cell-autonomous yet orchestrated by a master pacemaker in the hypothalamus ^7–10^. Cellular circadian clocks are primarily driven by coupled negative feedback loops formed between a Bmal1/Clock heterodimer and its transcriptional targets: cryptochrome (Cry), period (Per), and Rev-erb, which in turn inhibit Bmal1/Clock ^11^. In proliferating cells, several circadian proteins crosstalk to cell cycle proteins ^12–15^. For example, Bmal1 represses the expression of Myc ^16, 17^, a transcription factor that promotes cell proliferation; Rev-erb represses the expression of p21 ^Cip1^ (p21 for short) ^18^, a cyclin-dependent kinase (Cdk) inhibitor of G1 and S Cdks (Cdk4,6 and Cdk2); Bmal1/Clock activates Wee1 ^19^, leading to the suppression of Cyclin B1/Cdk1 activity that is critical to mitosis, while Cry does the opposite by inhibiting Wee1 ^19^. These molecular interactions lead to the circadian regulation of the proliferative cell cycle ^17–20^. However, how the circadian clock regulates the transition to proliferation from quiescence remains largely unknown.

Cellular quiescence is not a homogeneous state but heterogeneous ^1, 21, 22^. The likelihood of quiescence-to-proliferation transition is reversely correlated with quiescence depth. Upon growth stimulation, cells at deeper quiescence are less likely to, and take longer time if they do, reenter the cell cycle and initiate DNA replication than cells at shallower quiescence ^21, 23, 24^. Deeper quiescence is often observed in aging cells in the body ^25, 26^ or in cells remaining quiescent for longer durations in culture ^21, 27^. Shallower quiescence is seen in stem cells responding to tissue injury ^28, 29^ or related systemic signals ^30, 31^.

We have shown recently that quiescence depth is regulated by an Rb-E2f bistable switch and its interacting pathways ^21, 32, 33^. E2f transcription activators (E2f1-3a, referred to as E2f for short), by transactivating genes necessary for DNA synthesis and cell cycle progression, are both necessary and sufficient for cell cycle entry from quiescence ^34, 35^. E2f is repressed by Rb family proteins (referred to as Rb for short) in quiescent cells and activated upon stimulation with serum growth factors. E2f activation upon serum stimulation is mediated by multiple positive feedbacks leading to the phosphorylation and inhibition of Rb by Cyclin D (CycD)/Cdk4,6 and Cyclin E (CycE)/Cdk2 complexes. These integrated positive feedbacks establish the bistability at the Rb-E2f gene network level, converting graded and transient growth signals into an all-or-none E2f activation ^32, 36^, which further triggers and couples with APC/C^CDH1^ inactivation via EMI1 and CycE/Cdk2, leading to an irreversible entry of the S-phase of the cell cycle and thus the quiescence-to-proliferation transition ^37, 38^. The serum threshold (i.e., minimum serum concentration) that activates the Rb-E2f bistable switch, the E2f-activation threshold for short, has been shown to determine quiescence depth ^21^.

In this study, we examined the effects of two circadian proteins, Cry and Rev-erb, on the cellular transition from quiescence to proliferation. We anticipated that Cry and Rev-erb activities, via their known roles in upregulating Myc and downregulating p21, respectively, might lead to shallower quiescence by reducing the E2f-activation threshold ^21^. However, we observed the opposite in our experiments. Through a comprehensive modeling search and follow-up experiments, we found both Cry and Rev-erb play novel roles that converge to downregulate CycD/Cdk4,6 activity, leading to an ultrasensitive increase of the E2f-activation threshold and quiescence depth.

## RESULTS

### Circadian protein Cry deepens cellular quiescence

Earlier studies have suggested that Bmal1 inhibits Myc expression, and consistently, Cry upregulates Myc ^16, 17^. As Myc promotes E2f activation ^39^, we tested the potential effect of Cry on the quiescence-to-proliferation transition. We started by applying a recently developed specific Cry agonist KL001 ^40^ to rat embryonic fibroblasts (REF/E23 cells). When treated with KL001 (≤ 40 μM, below its cytotoxicity level, Fig S1A), REF/E23 cells induced to quiescence by serum starvation did exhibit a modest but statistically significant increase of Myc protein (Fig. S2A); this increase, however, became insignificant when quiescent REF/E23 cells were stimulated to enter the cell cycle (10 and 14 hours after serum stimulation, Fig. S2B). Assuming the modest increase of Myc in quiescent cells might facilitate their E2f activation and transition into proliferation, we expected quiescence depth might be slightly reduced, if any, under KL001 treatment. To our surprise, we found KL001 treatment deepened quiescence: with increasing KL001 doses, increasing serum concentrations were needed to drive similar percentages of cells to reenter the cell cycle (arrow pointed, ~45%; Fig. 1A, based on EdU incorporation, EdU+; Fig. S2C, based on the “On”-state of an E2f-GFP reporter, E2f-ON ^32^). Consistently, when stimulated at a given serum concentration (e.g., 8% serum, Fig. 1A and S2C), the percentage of cells that reentered the cell cycle decreased with increasing KL001 doses.

**Figure 1.**
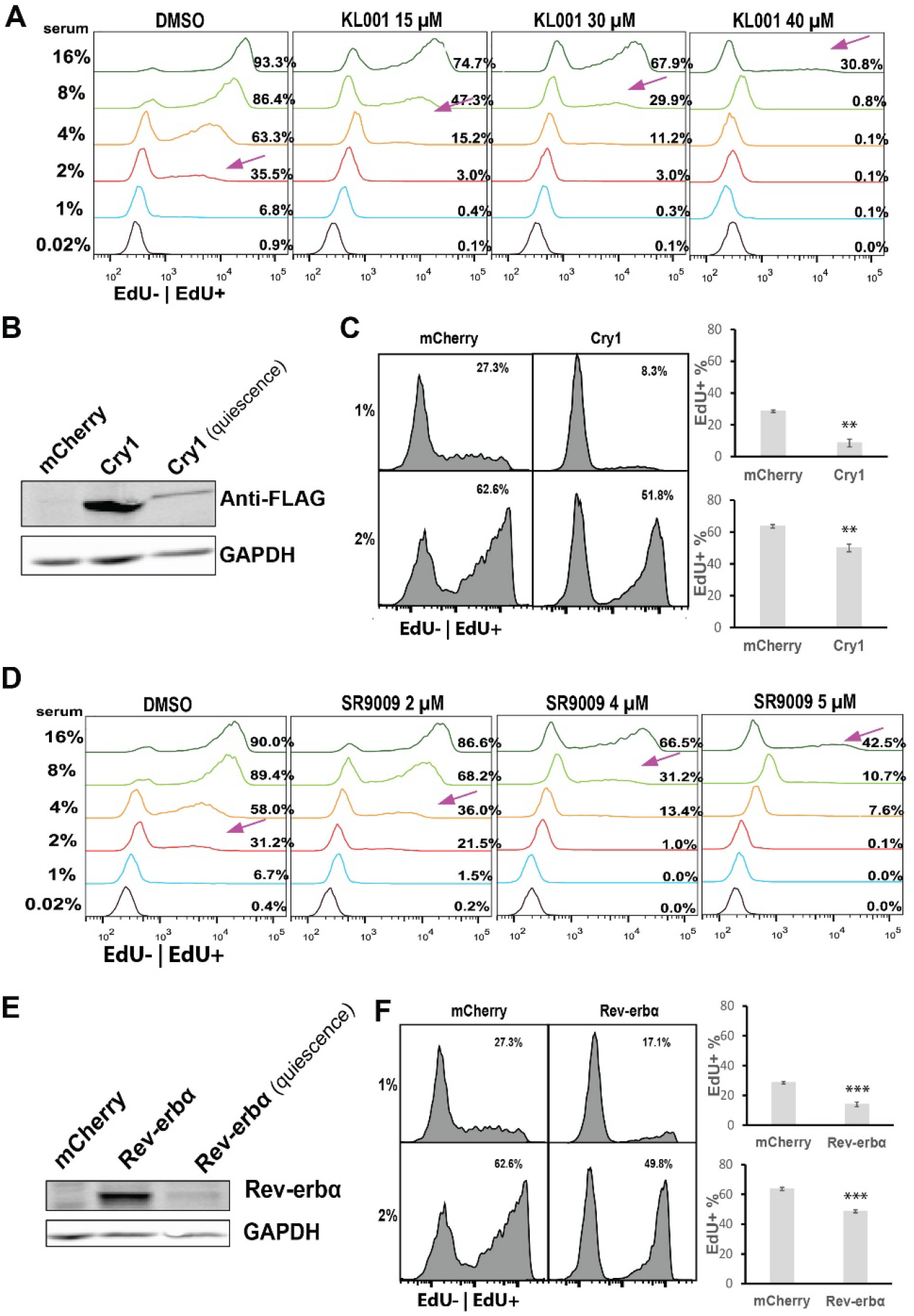
Cry and Rev-erb drive cells to deeper quiescence. (A) Effect of Cry agonist KL001 on quiescence depth. REF/E23 cells were first induced to quiescence by serum starvation for 2 days, then treated with the agonist at the indicated concentrations in starvation medium for 1 day; cells were subsequently stimulated by switching to growth medium containing serum and agonist at the indicated concentrations for the indicated durations. This protocol, serum stimulation (STI) of 3-day quiescent cells under agonist treatment (STI.3dq/agonist for short), was used for all agonist-related tests in this study unless otherwise noted. Indicated to the right of individual histograms are the percentages of cells becoming EdU+ after 24 hours of serum stimulation. Arrows indicate the approximate serum concentrations resulting in EdU+% = 45%. (B) Ectopic Cry1 expression. Proliferating cells were transfected with a FLAG-tagged Cry1 vector or a mCherry control and then subjected to Western blot with a FLAG antibody. “Quiescence” indicates the Western blot performed in Cry1-transfected cells induced to quiescence by 2-day serum-starvation. (C) Effect of ectopic Cry1 on quiescence depth. Cry1- or mCherry-transfected cells were induced to quiescence by 2-day serum-starvation, stimulated with serum at the indicated concentrations for 30 hours, and assayed for EdU+%. Error bar, SEM (n = 3). **p < 0.01, ***p<0.001 (1-tailed t-test; the same below). (D) Effect of Rev-erb agonist SR9009 on quiescence depth. Quiescent cells were serum stimulated following the STI.3dq/agonist protocol. Serum and SR9009 concentrations are as indicated. EdU+% at the 24-hour after serum stimulation are shown to the right of individual histograms. Arrows indicate the approximate serum concentrations leading to EdU+% = 45%. (E) Ectopic Rev-erbα expression. Proliferating cells were transfected with a Rev-erbα vector or a mCherry control and then subjected to Western blot with a Rev-erb antibody. “Quiescence” indicates the Western blot performed in Rev-erb-transfected cells induced to quiescence by 2-day serum-starvation. (F) Effect of ectopic Rev-erb on quiescence depth. Rev-erb- or mCherry-transfected cells were induced to quiescence by 2-day serum-starvation, stimulated with serum at the indicated concentrations for 30 hours, and assayed for EdU+%. Error bar, SEM (n = 3). **p < 0.01.

Similarly, quiescence deepened with the treatment of a different Cry agonist or with ectopic Cry1 expression. First, when REF/E23 cells were treated with the 2^nd^ Cry agonist, KL002, increasing serum concentrations were needed to drive similar percentages of cells to reenter the cell cycle (Fig. S2D, based on EdU+%; Fig. S2E, based on E2f-ON%). Consistently, when stimulated at a given serum concentration, the percentage of cells that reentered the cell cycle decreased with increasing KL002 doses (Fig. S2 D and E). Second, in quiescent REF/E23 cells transfected with a Cry1 vector and expressing ectopic Cry1 (albeit at a much lower level in quiescence than in growing condition, Fig. 1B), the percentage of cells that reentered the cell cycle (EdU+%) in response to serum stimulation decreased (p < 0.01) compared to that in the mCherry-transfection control (driven by the same CMV promoter, Fig. 1C). Our results from two Cry agonists and ectopic Cry expression, put together, suggested that Cry drove cells to deeper quiescence, instead of facilitating the quiescence-to-proliferation transition.

### Circadian protein Rev-erb deepens cellular quiescence

Next, we examined another link between circadian proteins and the Rb-E2f switch: Rev-erb inhibits the expression of Cdk inhibitor (CKI) p21 ^18^, while p21 was known to deepen quiescence by increasing the E2f-activation threshold ^21^. Therefore, we expected Rev-erb to reduce the E2f-activation threshold and thus quiescence depth. When we treated REF/E23 cells with a Rev-erb agonist SR9009 (≤ 5 μM, below its cytotoxicity level, Fig S1B), we did observe a significant decrease of p21 protein level at intermediate and high SR9009 doses (4 and 5 μM) both in quiescence (Fig. S3A) and during cell cycle entry (10 and 14 hours after serum stimulation, Fig. S3B). However, SR9009-treated cells did not move to shallower quiescence but deeper: with increasing SR9009 doses, increasing serum concentrations were needed to drive similar percentages of cells to reenter the cell cycle (arrow pointed, ~45%; Fig. 1D, based on EdU+%; Fig. S3C, based on E2f-ON%). When stimulated with serum at a given concentration, the percentage of cells that reentered the cell cycle decreased with increasing SR9009 doses (Fig. 1D and Fig. S3C). Similarly, treating cells with the 2^nd^ Rev-erb agonist, SR9011, also deepened quiescence (Fig. S3D and E). Consistent with the effects of Rev-erb agonists, ectopic Rev-erb expression deepened quiescence: in quiescent REF/E23 cells transfected with a Rev-erb vector and expressing ectopic Rev-erb (albeit at a much lower level in quiescence than in growing condition, Fig. 1E), the percentage of cells that reentered the cell cycle (EdU+%) in response to serum stimulation decreased (p < 0.01) compared to that in the mCherry-transfection control (driven by the same CMV promoter, Fig. 1F). Combining our results based on two agonists and ectopic expression, it suggested that Rev-erb drove cells to deeper quiescence instead of facilitating the quiescence-to-proliferation transition.

### Both Cry and Rev-erb downregulate CycD/Cdk4,6 to deepen quiescence as predicted by exploratory model search

Circadian proteins Cry and Rev-erb both deepened quiescence unexpectedly, indicating certain mechanistic links were missing in our understanding between these circadian proteins and cellular quiescence. As our earlier studies showed that the E2f-activation threshold determines quiescence depth ^21, 36^, this threshold mechanism provides a likely target for circadian regulation. We therefore searched for the potential missing links connecting Cry and Rev-erb to the E2f-activation threshold.

The Rb-E2f gene network is a complex system comprised of over 90 gene nodes involved in intertwined transcriptional controls and signaling cascades ^41–45^. It would be time- and labor-intensive to test the candidate links one by one in experiments without first effectively narrowing down the candidates. To this end, we took advantage of our previously established mathematical model of the Rb-E2f bistable switch ^32^ and used computer simulation to predict the most likely missing link(s) responsible for the quiescence-deepening effects of Cry and Rev-erb. Our previous Rb-E2f bistable switch model considered five coarse-grained network modules: Myc, CycD/Cdk4,6, Rb, E2f, and CycE/Cdk2 (Fig. 2A). Upon serum growth signals, Myc and CycD/Cdk4,6 are upregulated. Myc promotes E2f expression; CycD/Cdk4,6 phosphorylates Rb and partially de-represses E2f. E2f activates CycE/Cdk2, which further phosphorylates Rb and de-repress E2f, forming a positive feedback loop. E2f transactivates its own expression, forming another positive feedback loop that reinforces E2f activation. Next, we considered Cry or Rev-erb (the C/R module, Fig. 2A) might directly or indirectly interact with any or all of the five modules (i.e., five possible links), and exert one of three net effects (up-regulation, down-regulation, or no effect). For example, the two literature links we started the study with, Cry upregulating Myc and Rev-erb inhibiting p21, were reflected in C/R upregulating Myc and Cyclin/Cdk modules (CycD/Cdk4,6 and CycE/Cdk2, via downregulating p21), respectively. Considering the possible combinations of 5 links with 3 effects each, 3^5^ = 243 different topologies could be generated to connect C/R to the Rb-E2f switch.

**Figure 2.**
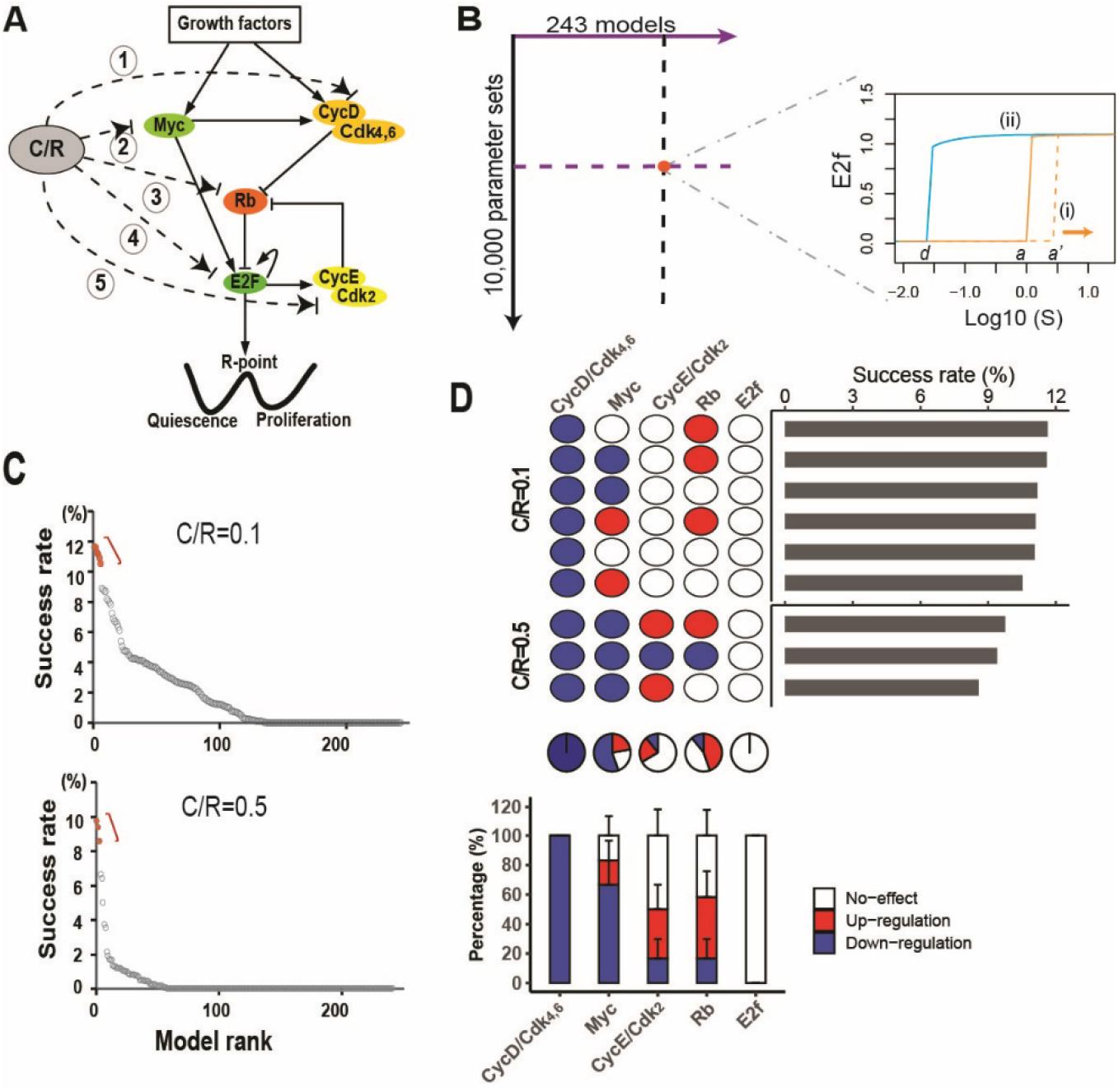
Modeling search for the missing links of how Cry and Rev-erb deepen quiescence. (A) Cry or Rev-erb (C/R) may crosstalk with any or all of the five Rb-E2f network modules, and each of the five links (1-5) can have one of the three possible net effects: upregulation, downregulation, and no-effect, thus generating 3^5^ = 243 possible network topologies. (B) Model search and simulation. Each of the 243 models was simulated with 10,000 random parameter sets; with each parameter set, the model was evaluated according to two criteria: (i) E2f-activation threshold *a*’ ≥ 3 (serum units); and (ii) bistability (as in the base model, E2f-activation threshold *a* > E2f-deactivation threshold *d*). S (x-axis), serum unit; E2f (y-axis), steady-state E2f level. Solid orange and blue curves indicate E2f serum-responses in the base model simulated from the quiescence and proliferation initial conditions, respectively. For simplicity, only the E2f serum-response from quiescence simulated with one random parameter set (dashed orange curve) is shown. See Materials and methods for details. (C) Model ranking. The 243 models were ranked from left to right (x-axis) based on model success rate (y-axis), which indicated the percentage of events (random parameter sets) in which the model simulation outcome fulfilled the two criteria (i) and (ii). Simulation results with the C/R input level of 0.1 and 0.5 are shown at the top and bottom, respectively. (D) Link features of top-ranked models. (Top) Highest-ranked models with similar success rates at C/R = 0.1 and 0.5, respectively (red dots in C). The link features in each model are shown according to the upregulation (red), downregulation (blue), or no-effect (white) of the indicated target node by C/R. (Middle) Pie chart of the percentage of each link feature at the indicated target node among the combined 9 models (top). (Bottom) The average percentage of each link feature at the indicated target node between the two model groups (C/R = 0.1 and 0.5, respectively, top). Error bar, SD.

We constructed a library of 243 ordinary differential equation (ODE) models to represent the 243 network topologies. Based on our experimental observations, we set two search criteria for the most likely missing link(s) between C/R and the Rb-E2f switch: (i) increasing the E2f-activation threshold, and (ii) maintaining the network bistability. Specifically, for criterion i, we set the E2f-activation threshold to increase from 1.0 in the previous base model ^32^ to ≥ 3.0, since comparable EdU+% was obtained with 1% serum in the DMSO control and ~3% serum under KL001 and SR9009 treatments at intermediate doses (KL001, 30 μM; SR9009, 4 μM; Fig. 1A and D). Criterion ii was set because we expected the Rb-E2f bistable switch to remain critical to the proper quiescence-to-proliferation transition under circadian regulation; it was also consistent with the bimodal E2f expression observed under KL001 and SR9009 treatments (similar to the DMSO control, Fig. S2C and S3C).

We subsequently simulated each of the 243 ODE models with a collection of random parameter sets. Each set contained 5 model parameters that dictated the strengths of the 5 links from C/R to Rb-E2f network modules (*I_1-5_*, Tables S1 and S2), with parameter values simultaneously and randomly selected within the numerical ranges as determined in our previous modeling studies ^32, 46^. We then calculated the success rate of each of the 243 models in fulfilling the search criteria (i) and (ii) across the random parameter sets (Fig. 2B, see Materials and Methods for detail). In this regard, we also applied two different C/R input levels to account for relatively low and high agonist doses, respectively (Fig. 2C). The models with the highest success rates (i.e., most robust against parameter variations ^46–48^) were considered the most likely explanations for how Cry and Rev-erb increased the E2f-activation threshold and deepened quiescence as we observed in experiments (Fig. 1A and D).

We found the compositions of the top 10 models remained the same after 5,000 random parameter sets at C/R = 0.1 (Table S3) and 3,750 parameter sets at C/R = 0.5 (Table S4), respectively, suggesting the model search results were stabilized. As seen in Fig. 2C, the final model success rates (after 10,000 parameter sets) declined rapidly moving away from the top-ranked models at each C/R input level, suggesting that a limited number of model topologies were viable for the C/R activity to deepen quiescence. That said, no single model “winner” stood out. For example, the top 6 models at C/R = 0.1 (red dots, Fig. 2C top panel) formed a cluster; within the cluster, any two neighboring models A and B exhibited similar success rates (s.r.), with the relative s.r. difference (s.r._a_-s.r._b_)/s.r._a_ < 10%. The same was true for the top 3 models at C/R = 0.5 (red dots, Fig 2C bottom). Comparing these top s.r. models (6 at C/R = 0.1; 3 at C/R = 0.5), one uniquely shared feature became apparent: the downregulation of CycD/Cdk4,6 by C/R in every model (Fig. 2D). Alternatively, when we chose the 10 top-ranked models at each C/R input level (10 at C/R = 0.1; 10 at C/R = 0.5) as another high-s.r. model selection approach, CycD/Cdk4,6 downregulation by C/R was again the number one shared feature (Fig. S4). These model simulation results predicted that Cry and Rev-erb likely induced deep quiescence by primarily targeting and downregulating CycD/Cdk4,6 activity. We will further interpret these modeling results in Discussion.

### Experimental support for CycD/Cdk4,6 as the primary target of Cry and Rev-erb

To test our model predictions, we measured the responses of CycD/Cdk4,6 complex components to Cry and Rev-erb and compared them to those of CycE/Cdk2 complex components. Specifically, following the same STI.3dq/agonist protocol (Fig. 1), we treated REF/E23 cells with Cry and Rev-erb agonists and measured the protein levels of D-type cyclins CycD1 and CycD3 (CycD2 is not expressed in REF/E23 cells ^23^), Cdk4, and Cdk6, as well E-type cyclins (CycE1 and CycE2) and Cdk2 in quiescence and during cell cycle entry upon serum stimulation (Fig. 3).

**Figure 3.**
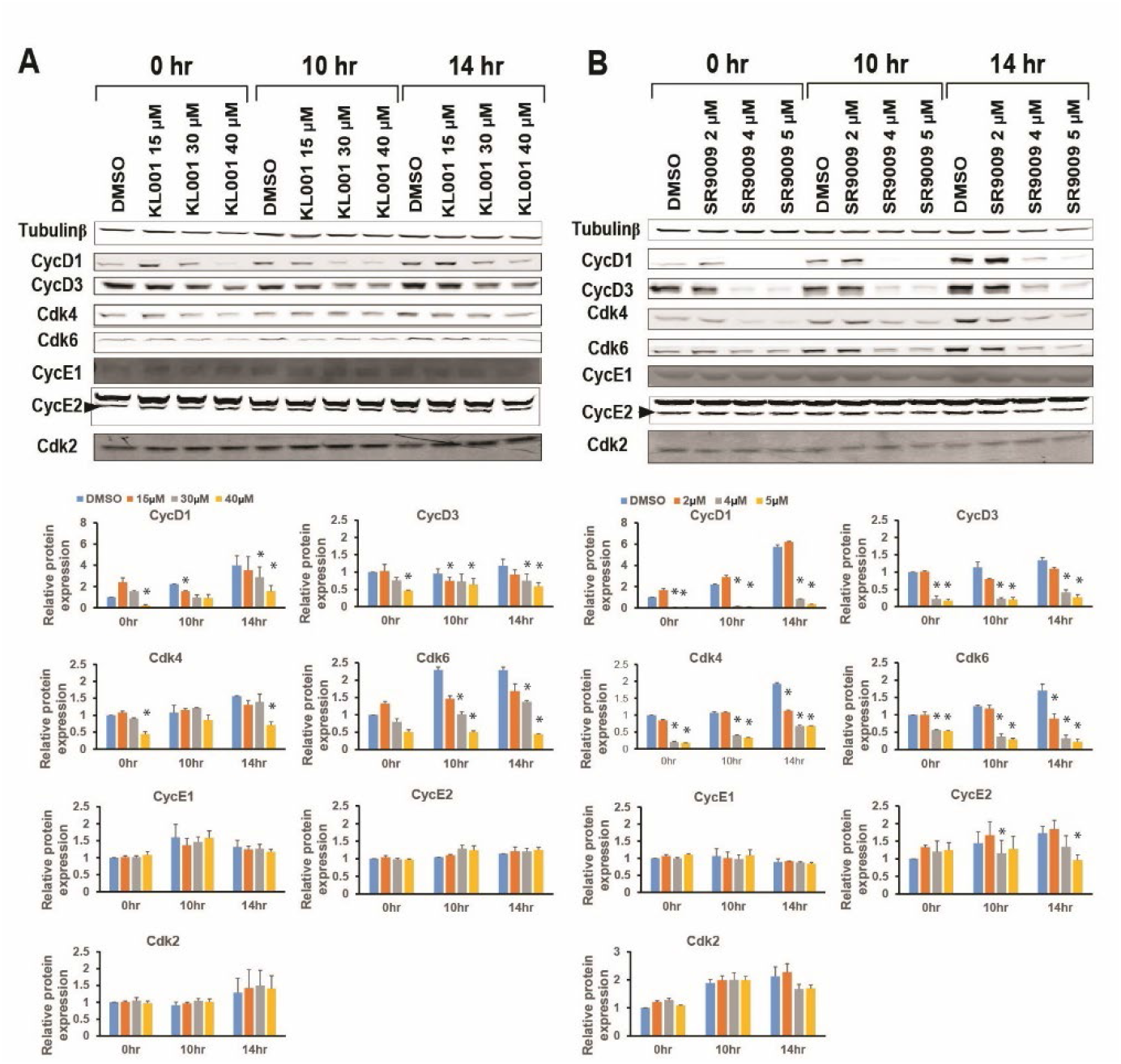
Cry and Rev-erb downregulate protein components of CycD/Cdk4,6 but not CycE/Cdk2. The responses of individual protein components of CycD/Cdk4,6 and CycE/Cdk2 to the Cry agonist KL001 (A) and Rev-erb agonist SR9009 (B) at the indicated doses were measured following the STI.3dq/agonist protocol. Protein levels were measured by Western blot in quiescence (0 hr) and during cell cycle entry (10 and 14 hr after stimulated with 4% serum) and normalized to the 0-hr DMSO control. Error bar, SEM (n=2); *p < 0.05 (1-tailed t-test).

We observed a significant downregulation of each tested CycD/Cdk4,6 complex component in response to both the Cry agonist KL001 (Fig. 3A) and Rev-erb agonist SR9009 (Fig. 3B). This general pattern occurred across the three tested time points (0, 10, and 14 hours upon serum stimulation), especially at the medium and high agonist doses (KL001, 30 and 40 μM; SR9009, 4 and 5 μM; see Discussion for the low dose). This pattern of CycD/Cdk4,6 downregulation in response to Cry and Rev-erb agonists was in stark contrast to that of CycE/Cdk2 complex components, which exhibited non-significant changes overall (Fig. 3). These experimental observations were in good agreement with our model simulation results (Fig. 2D and S4), showing a convergent downregulation of CycD/Cdk4,6 by Cry and Rev-erb.

Similarly, we measured the responses of other proteins (Rb; Rb phosphatases PP1 and PP2A; Myc; CKIs p21, p27, and p16) in the Rb-E2f bistable switch network to Cry and Rev-erb agonists (Fig. S5). None of the observed responses, if any, would explain quiescence deepening (see Discussion). Put together, our experimental results supported the model-predicted unique role of CycD/Cdk4,6 as the convergent target of the circadian regulation by Cry and Rev-erb on quiescence depth.

## DISCUSSION

The circadian clock aligns diverse cellular functions to periodic daily environmental changes. In this study, we investigated the effects of two key circadian proteins, Cry and Rev-erb, on cellular quiescence. We found upregulating Cry and Rev-erb drove cells into deeper quiescence, opposite to what we had hypothesized based on literature. To identify the missing links in our understanding, we evaluated an assembly of potential network models and tested the converged predictions of the top-ranked models experimentally. Our results suggested that both Cry and Rev-erb deepen quiescence by primarily downregulating the CycD/Cdk4,6 complex components in the bistable Rb-E2f gene network.

Why does the CycD/Cdk4,6 module play a unique role, targeted by both Cry and Rev-erb, in mediating quiescence deepening? To move into deep quiescence, a cell needs to increase its E2f-activation threshold while maintaining the Rb-E2f bistable switch for the proper quiescence-proliferation transition. In this regard, altering the activities of different Rb-E2f network modules has different consequences, depending on their positions and roles in the network (Fig. 2A). For example, changing Cyclin/Cdk activity has a stronger effect than changing Rb synthesis in increasing the E2f-activation threshold (determined by the ratio of unphosphorylated Rb over free E2f) ^21^. Between the two G1 Cyclin/Cdks, late G1 CycE/Cdk2 hyper-phosphorylates Rb, and CycE/Cdk2 downregulation is associated with cell cycle arrest or exit into quiescence in proliferating cells ^49, 50^. To drive quiescent cells deeper by targeting CycE/Cdk2, however, can be problematic. This is because downregulating CycE/Cdk2 weakens the mutual-inhibition (i.e., positive feedback) loop between Rb and E2f that is essential to the network bistability ^46^ and consequentially the proper quiescence-to-proliferation transition. Similarly, targeting and inhibiting E2f to increase the E2f-activation threshold could be problematic, as repressing E2f weakens both positive feedbacks (Rb-E2f mutual-inhibition and E2f auto-activation) in the Rb-E2f network underlying the network bistability. By contrast, CycD/Cdk4,6 is upstream of and not directly involved in the positive feedback loops between Rb and E2F (Fig. 2A). We therefore expect that targeting to inhibit CycD/Cdk4,6 can better divide and conquer the needs to both increase the E2f-activation threshold and maintain the Rb-E2f bistable switch. Consistently, Cdk6 expression level was found to regulate the quiescence depth of human hematopoietic stem cells and impact their long-term preservation ^51^.

Relatedly, in proliferating cells, CycD level appears to reflect the protein synthesis rate of the mother cell, and the CycD level inherited from the mother cell is a key determinant of the proliferation-quiescence bifurcation of daughter cells ^52, 53^. On another note, a recent study suggested that CycD/Cdk4,6 mono-phosphorylates but does not inhibit Rb, and it meanwhile activates CycE/Cdk2 via an unidentified mechanism ^54^. Although this new finding and the classic model differ in whether CycD/Cdk4,6 directly inhibits Rb, they are consistent in the role of CycD/Cdk4,6 in leading to CycE/Cdk2 activation and initiating the positive mutual-inhibition loop between Rb and E2f, which eventually leads to E2f activation and the passage of the restriction point during the quiescence-to-proliferation transition ^55–60^.

A closer look of CycD/Cdk4,6 responses to Cry and Rev-erb agonists showed that the protein levels of CycD1, CycD3, Cdk4, and Cdk6 decreased noticeably at the medium and high doses of KL001 and SR9009, but not at their low doses (15 μM KL001, Fig. 3A; 2 μM SR9009, Fig. 3B). How would this result at low dose conditions explain the then still significantly deepened quiescence (Fig. 1A and D)? It turns out Rb phosphorylation can be ultrasensitive to CycD/Cdk4,6 activity. That is, a small downregulation of CycD/Cdk4,6 may cause a much larger decrease of Rb phosphorylation during cell cycle entry (see the 10- and 14-hr sim curves, Fig. 4A), as predicted by our Rb-E2f bistable switch model reflecting the phosphorylation-dephosphorylation zero-order ultrasensitivity ^61^. As a rough estimate, CycD/Cdk4,6 activity under the low doses of KL001 and SR9009 was reduced by about 30% during cell cycle entry (considering joint cyclin, Cdk, and CKI levels, Table S5), which could result in an over 80% reduction of Rb phosphorylation based on model simulations (Fig. 4A). We note that the estimate of CycD/Cdk4,6 was based on several assumptions (Table S5) and may not be accurate. That said, our experimental observations of the phospho-Rb level (S807/S811) across varying KL001 and SR9009 doses (low, medium, and high, Fig. 4 B and C) were in good agreement with the model predictions considering the estimated CycD/Cdk4,6 activities (Fig. 4A). This ultrasensitive decrease of Rb phosphorylation increased the E2f-activation threshold and thus deepened quiescence in our model, leading to noticeably fewer cells able to reenter the cell cycle upon serum stimulation (Fig. 4D).

**Figure 4.**
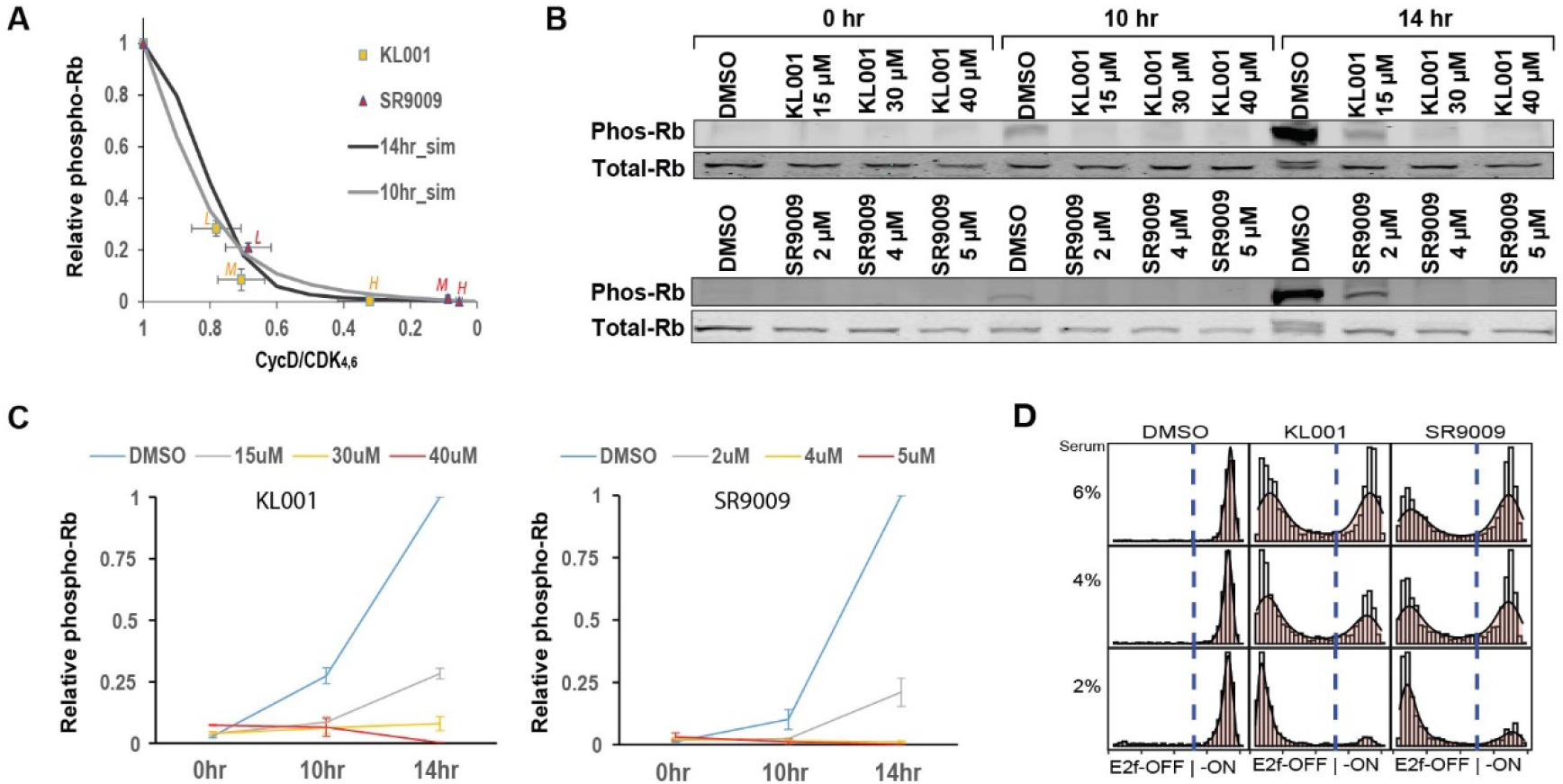
Ultrasensitive effects of CycD/Cdk4,6 on Rb phosphorylation and quiescence depth. (A) Relationship between CycD/Cdk4,6 (x-axis) and relative phospho-Rb = (phosphorylated Rb)/(total Rb) (y-axis). Model simulated results at the 10-hr (gray curve) and 14-hr (black curve) after serum stimulation are shown together with experimentally estimated data points (CycD/Cdk4,6, from Table S5; relative phospho-Rb, from C) under the treatments of KL001 (yellow squares) and SR9009 (red triangles) at the low (L), medium (M), and high (H) doses, respectively. (B) Effects of Cry and Rev-erb agonists (KL001, top; SR9009, bottom) on Rb phosphorylation. Quiescent cells were serum (4%) stimulated following the STI.3dq/agonist protocol in the presence of agonists at the indicated concentrations. The levels of phosphorylated Rb protein (S807/S811) were measured by Western blot at the 0-, 10-, and 14-hr time points after serum stimulation. (C) Quantification of the relative phospho-Rb in response to Cry and Rev-erb agonists (KL001, left; SR9009, right). Levels of phosphorylated Rb and total Rb were quantified from Western blots as in B. The relative phospho-Rb value of the 14-hr DMSO control is set to 1.0. Error bar, SD. (D) Stochastic simulations of quiescence exit under the low doses of KL001 (15 μM) and SR9009 (2 μM). In each panel with the indicated condition, 1000 cells were simulated according to the relative Phospho-Rb level (14 hr) as in B, and the distribution of simulated E2f molecule numbers at the 24-hr after stimulation (4% serum) are shown (x-axis).

CycD/Cdk4,6 activity can be reduced by either downregulating CycD and Cdk4,6 or upregulating Cdk inhibitors (CKIs), including Cip/Kip proteins (most notably p21 and p27) and INK4 proteins (most notably p16). In REF/E23 cells treated with Cry and Rev-erb agonists, the p16 level did not change significantly while the levels of p21 and p27 mostly decreased but not increased (Fig. S5). These observations suggest that CKIs are not responsible for CycD/Cdk4,6 downregulation by Cry and Rev-erb. Since p16 is a marker of senescent cells, that it is not targeted and upregulated by circadian proteins in quiescent cells was anticipated. Yet, p21 and p27 could be viable options since they have been shown to drive deep quiescence in different contexts ^21, 62^. We suspect that Cry and Rev-erb do not upregulate p21 and p27 to deepen quiescence is likely due to the specific regulatory mechanisms they employ to modulate the Rb-E2f bistable switch (see below).

Our modeling and experimental study here identified two novel connections from circadian proteins Cry and Rev-erb converging to downregulate CycD/Cdk4,6. The natures of these two newly discovered links remain unknown, such as how Cry and Rev-erb simultaneously downregulate multiple components (CycD1, CycD3, Cdk4 and Cdk6), and whether such regulations are direct or indirect. Similarly, we observed that both Cry and Rev-erb agonists reduced total Rb protein level (which would not deepen quiescence) but did not change the levels of Rb phosphatases PP1 and PP2A (Fig. S5). How both Cry and Rev-erb converge to similar regulatory patterns targeting a common subset of Rb-E2f network components are interesting questions that we hope to address in future studies.

As circadian proteins, Cry and Rev-erb levels fluctuate diurnally. Given upregulating Cry and Rev-erb deepened quiescence as observed in this study, we speculated that cellular quiescence depth might fluctuate diurnally too. Indeed, this is what we observed: circadian changes in the rate of quiescence-to-proliferation transition in REF/E23 cells upon growth stimulation (Fig. S6). Further studies are needed to test and confirm which CycD/Cdk4,6 components fluctuate accordingly and are responsible for this phenomenon. Also, studies are needed to answer whether circadian fluctuation of quiescence depth occurs in various body tissues, resulting in different rates of tissue repair and regeneration at different times of the day. We also anticipate the differences of targeting CycD/Cdk4,6 and CycE/Cdk2 in regulating quiescence depth, as found in this study, may have implications in the applications of Cdk4,6 and Cdk2 inhibitors in clinical settings.

## MATERIALS AND METHODS

### Cell culture, quiescence induction and exit

Rat embryonic fibroblasts REF52/E23 cells stably harboring an E2f1 promoter-driven destabilized GFP reporter were derived previously as in ^32^. Cells were routinely passed at a sub-confluent level and cultured in Dulbecco’s Modified Eagle’s Medium (DMEM) (No. 31053, Gibco, Thermo Fisher) with 10% of bovine growth serum (BGS, No. SH30541, Hyclone, GE Healthcare). To induce quiescence, growing cells were seeded at around 10^5^ cells per well in 6-well culture plates (No. 353046, Corning Falcon), washed twice with DMEM, followed by culture in serum-starvation medium (0.02% BGS in DMEM) for 2 days or as indicated. To induce quiescence exit, serum-starvation medium was changed to serum-stimulation medium (DMEM containing serum at the indicated concentration), and cells were subsequently cultured for the indicated durations.

### Treatments of Cry and Rev-erb agonists

Cry agonists KL001 (No. 233624, EMD Millipore) and KL002 (No. 13879, Cayman) and Rev-erb agonists SR9009 (No. 554726, Sigma-Aldrich) and SR9011 (No. SML2067, EMD Millipore) were applied by being included in serum starvation and stimulation media at the indicated times and concentrations. DMSO was used as a vehicle control.

### E2f activity and quiescence exit (EdU incorporation) Assays

To measure E2f activity in individual cells, cells were harvested at the 24-hr time point after serum stimulation, fixed with 1% formaldehyde, and the fluorescence intensities of the E2f-GFP reporter in approximately 10,000 cells from each sample were measured using a LSR II flow cytometer (BD Bioscience). Flow cytometry data were analyzed using FlowJo software (v. 10.0). To assay for quiescence exit, EdU (2 μM) was added to serum-stimulation medium, and cells were collected at the indicated time points, followed by Click-iT EdU assay according to the manufacturer’s instruction.

### Ectopic expression

Growing REF/E23 cells were kept at sub-confluence and transfected with pfmh-hCry1 (a gift from Aziz Sancar; Addgene plasmid #25843) and pAdTrack-CMV FLAG Rev-erbα expression vectors (a gift from Bert Vogelstein; Addgene plasmid #16405) for ectopic expression of Cry1 and Rev-erbα, respectively, or with pCMV-mCherry (a gift from Lingchong You) as a control. Transfection was performed using Neon electroporation system (MPK5000, Invitrogen, Thermo Fisher) following the manufacturer’s protocol. Briefly, in each transfection, about 10^6^ cells were electroporated with 10 μg of plasmid DNA at 1800 volts with two 20-millisecond pulses in a 100 μl Neon tip.

### Western blotting

Cells were washed with DPBS once and then lysed in lysis buffer (50 mM Tris-HCl, pH 6.8, 2% sodium deoxycholate, 0.025% Bromophenol blue, 10% glycerol, 5%β-Mercaptoethanol). Extracted proteins were separated using SDS-PAGE and transferred onto nitrocellulose membranes. Immunoblot analyses were performed using antibodies against c-Myc (#sc-40), Rb (#sc-74562), Cyclin D1 (#sc-8396), Cyclin E2 (#sc-28351), Cdk2 (#sc-6248), Cdk4 (#sc-23896), p21 (#sc-53870), p27 (#sc-1641), PP1 (#sc-7482), and PP2A (#sc-13600) from Santa Cruz; Phospho-Rb ( S807/S811; #9308T), Cyclin D3 (#2936T), Cyclin E1 (#20808), and Cdk6 (#3136T) from Cell Signaling; p16 (#ab51243) from Abcam; Tubulin beta (#MAB3408) from EMD Millipore Corp; and GAPDH (#MA5-15738-D680) from Invitrogen.

### Model library generation and search

Regulatory effects of Cry or Rev-erb ([CR] in Table S1) on a node x in the Rb-E2f network were modeled by adding *m*\**w*\*[*CR*]/(*I*+[*CR*]) to the ODE *d[x]/dt* in our previously established Rb-E2f bistable switch model [19], with *m* = −1 (negative regulation), 0 (no regulation), or +1 (positive regulation); *w* = *k_x_* (matching the synthesis rate constant of x), and *I* being a random number uniformly distributed in the log range of 0.01~1.

Each of the 3^5^ = 243 models was simulated with the same set of 10,000 random parameter sets. With each parameter set, the activity of each node in the Rb-E2f network was simulated at 50 serum concentrations uniformly distributed in the log range between 0.01 and 20 (percent of serum, covering the conditions from serum starvation to saturation). To determine E2f bistability, at each serum concentration, the model with the initial condition (Table S1) corresponding to the quiescence state was simulated for 1000 model hours to reach the “switch-On” steady state. The “switch-On” steady-state values of individual nodes were then used as the initial condition of the proliferation state; the model was subsequently simulated for 1000 model hours to reach the “switch-Off” steady state. Simulation results were analyzed using an “in-house” Perl script, according to the criteria developed in our previous work ^46^ to determine E2f bistability. All simulations were performed using COPASI ^63^.

### Modeling phospho-Rb and CycD/Cdk4,6 downregulations

To simulate the CycD/Cdk4,6 downregulation under Cry and Rev-erb agonists, the CycD/Cdk4,6 term ([CD], Table S1) in the base model of the Rb-E2f bistable switch ^32^ was multiplied with a scaling factor (α = 0.1~1); phosphorylated Rb ([RP], Table S1) and total Rb ([R]+[RE]+[RP], Table S1) were accordingly derived by model simulation. Reversely, given a decrease in relative phospho-Rb under Cry and Rev-erb agonists as measured in the experiment, the scaling factor α of [CD] was derived by simulation and then applied to simulate the E2f serum response. In the time course simulation of the base model, the 7-hr model time aligned with the 14-hr experimental time according to phospho-Rb and E2f-GFP dynamics following serum stimulation. Accordingly, time in all model simulations was adjusted by 7 hours (e.g., the 17-hr model time was used to simulate serum responses at the 24-hr).

### Stochastic simulation

Similar to Ref. ^64, 65^, we built a Langevin-type stochastic differential equation (SDE) model based on the ODE model described in Table S1.

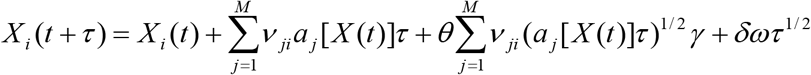

where *X*(*t*) = (*X*_1_(*t*), …, *X_n_*(*t*)) is the system state at time *t*. *X_i_*(*t*) is the molecule number of species *i* (*i* = 1, …, *n*) at time *t*. The time evolution of the system is measured based on the rates *a_j_*[*X*(*t*)](*j* = 1, …, *M*) with the corresponding change of molecule number *i* described in *ν_jl_*. Factors *γ* and *ω* are two independent and uncorrelated Gaussian noises. In this equation, the first two terms indicate deterministic kinetics, the third and fourth terms represent intrinsic and extrinsic noise, respectively. *θ* and *δ* are scaling factors to adjust the levels of intrinsic and extrinsic noise, respectively (*θ*=0.45, *δ*=30). Units of model parameters and species concentrations in the ODE model (Table S2) were converted to molecule numbers. We considered the cell reenters the cell cycle with the E2f-ON state, when the E2f molecule number at the 24^th^ hour after serum stimulation was larger than a cutoff value of 200. All SDEs were implemented and solved in Matlab.

## ACKNOWLEDGEMENTS

We thank Kotaro Fujimaki, Xiaojun Tian, and Tongli Zhang for critical readings and comments on the manuscript. This work was supported by grants from the NSF (#1463137 and 2034210 to GY) and NIH (GM-084905, a T32 fellowship to JSK) of the U.S.A., and the NSF of China (#31500676 to XW).

---

**Figure S1.**
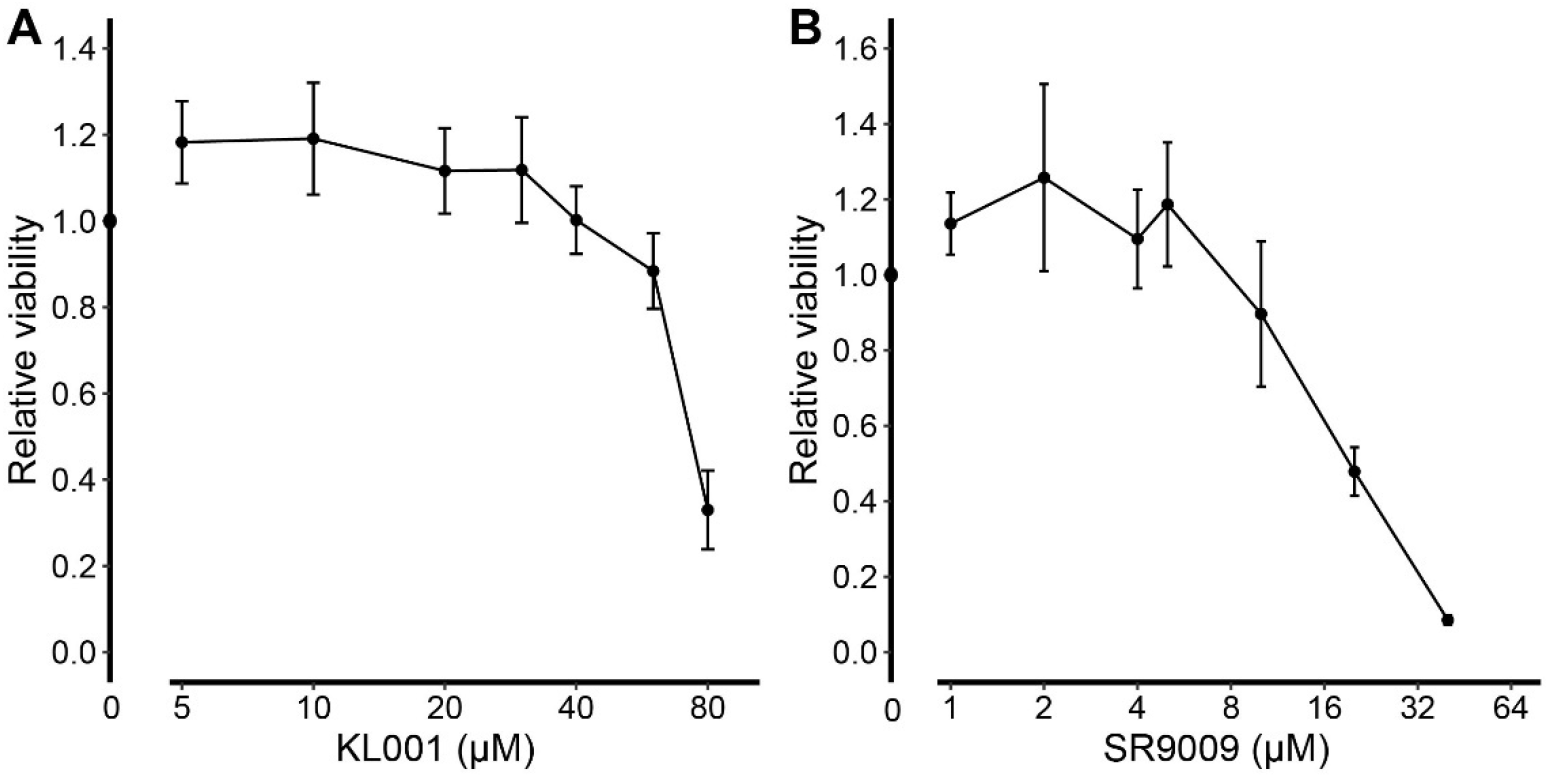
Cytotoxicity of Cry and Rev-erb agonists in REF52/E23 cells. Cells were treated with Cry agonist KL001 (A) and Rev-erb agonist SR90009 (B), respectively, at the indicated concentrations for 48 hours (concentration = 0 being the DMSO vehicle control). Relative viability (y-axis) refers to the ratio of the live cell count in a treated sample over that in the DMSO control. Live cell count was determined by the PI fluorescence assay as previously described ^66^. Error bar, SD (n = 3).

**Figure S2.**
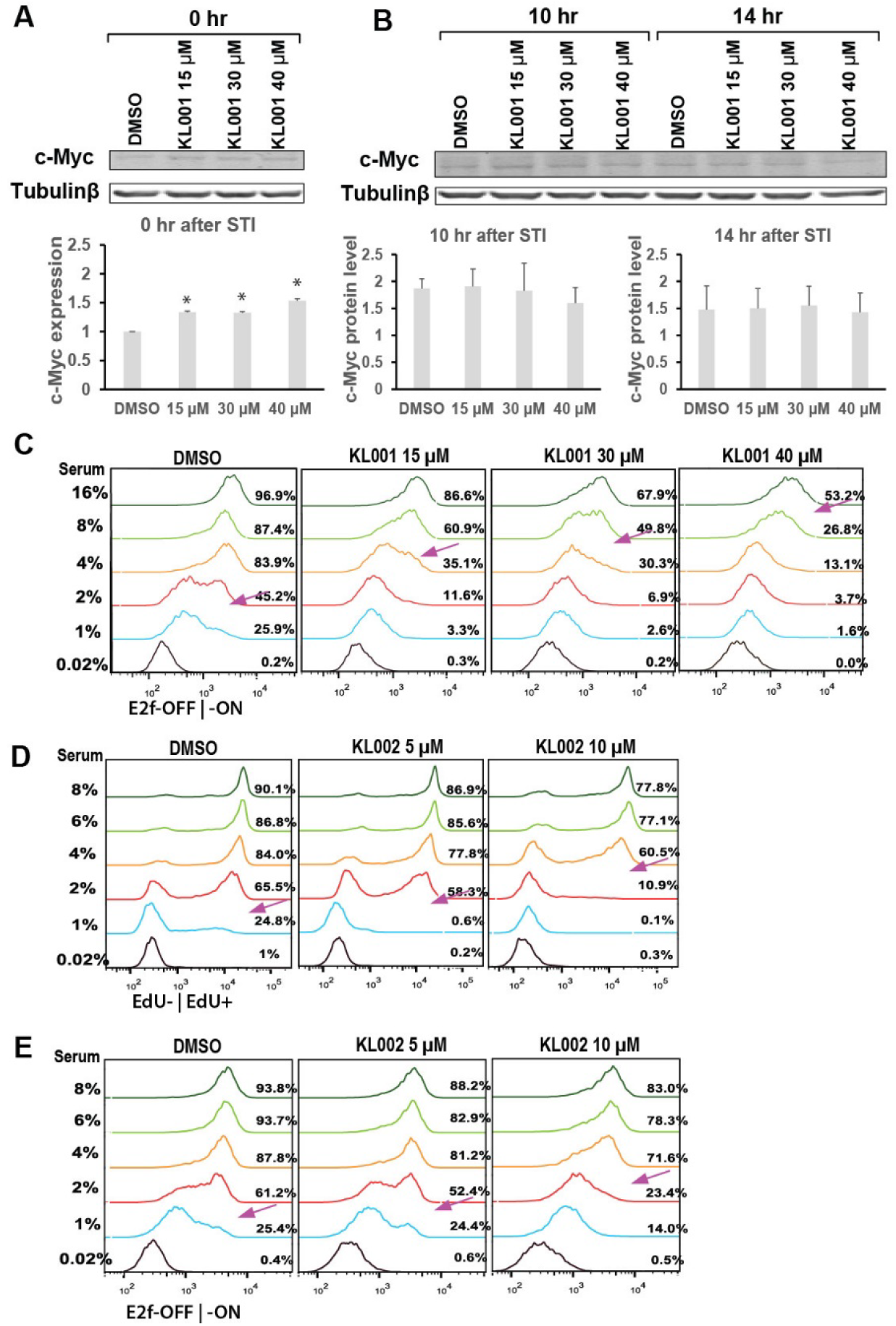
Cry agonists induce cells into deeper quiescence. (A-B) Effect of Cry agonist KL001 on Myc protein level. Quiescent cells were serum (4%) stimulated following the STI.3dq/agonist protocol in the presence of KL001 at the indicated concentrations. The protein levels of c-Myc were assessed by Western blot at (A) the 0 hr, i.e., in quiescent cells, and (B) the 10- and 14-hr time points after serum stimulation (STI). Error bar, SEM (n = 2). *p < 0.05. (C) Effect of KL001 on quiescence depth. Quiescent cells were serum stimulated following the STI.3dq/agonist protocol. Serum and KL001 concentrations are as indicated. The percentages of cells with E2f-ON at the 24-hr after serum stimulation are indicated to the right of individual histograms. Arrows indicate the approximate serum concentrations leading to E2f-ON% = 45%. (D-E) Effect of Cry agonist KL002 on quiescence depth. Quiescent cells were serum stimulated following the STI.3dq/agonist protocol. Serum and KL002 concentrations are as indicated. The EdU+% at the 30-hr (D) and the E2f-ON% at the 24-hr (E) after serum stimulation are indicated to the right of individual histograms. Arrows indicate the approximate serum concentrations leading to EdU+% (D) and E2f-ON% (E) = 45%, respectively.

**Figure S3.**
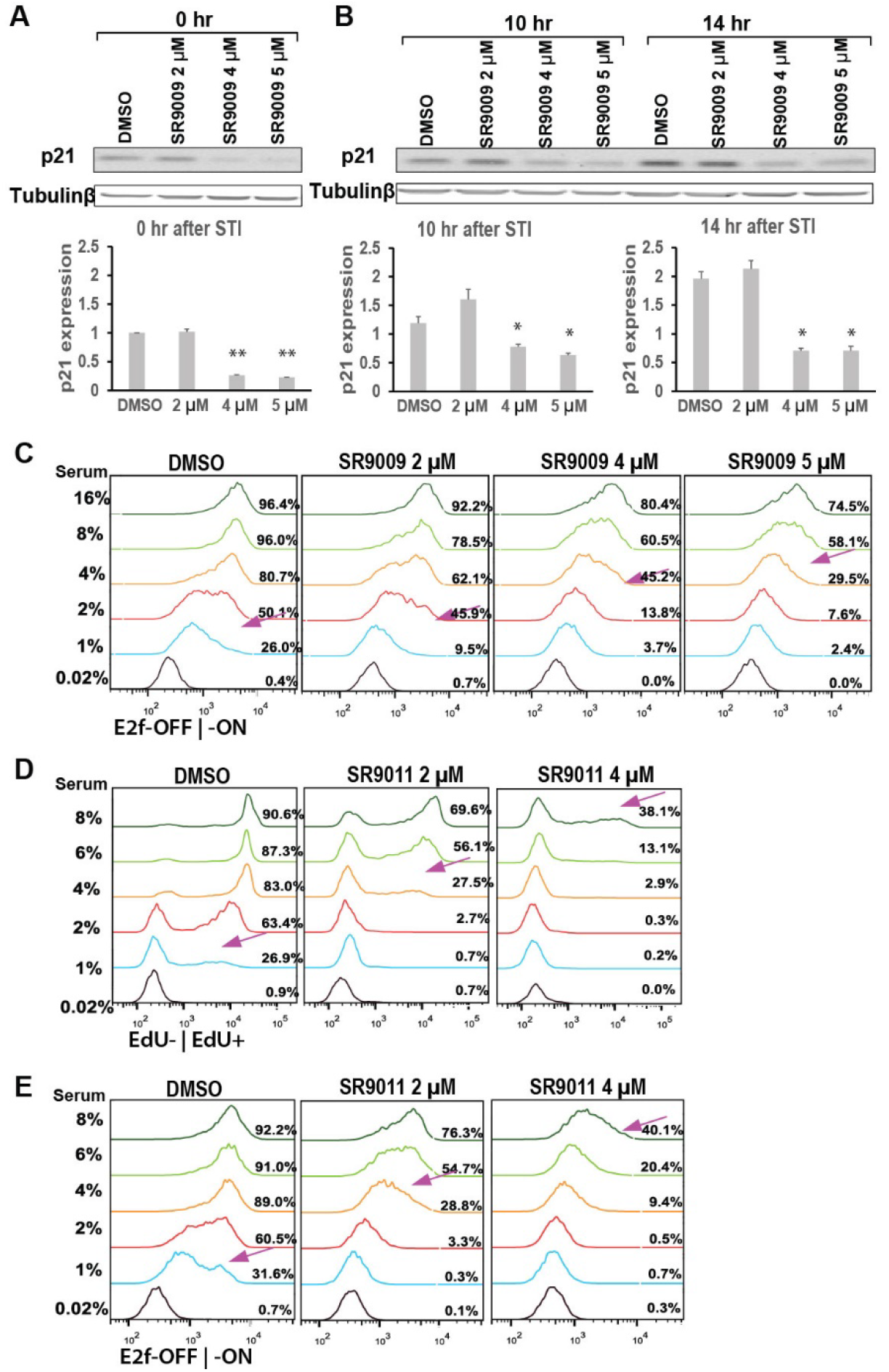
Rev-erb agonists induce cells into deeper quiescence. (A-B) Effect of Rev-erb agonist SR9009 on p21 protein level. Quiescent cells were serum (4%) stimulated following the STI.3dq/agonist protocol in the presence of SR9009 at the indicated concentrations. The protein levels of p21 were measured by Western blot at (A) the 0 hr (i.e., in quiescent cells), and (B) the 10- and 14-hr after serum stimulation (STI). Error bar, SEM (n = 2). *p< 0.05. (C) Effect of SR9009 on quiescence depth. Quiescent cells were serum stimulated following the STI.3dq/agonist protocol. Serum and SR9009 concentrations are as indicated. E2f-ON% at the 24-hr after serum stimulation are indicated to the right of individual histograms. Arrows indicate the approximate serum concentrations leading to E2f-ON% = 45%. (D-E) Effect of Rev-erb agonist SR9011 on quiescence depth. Quiescent cells were stimulated following the STI.3dq/agonist protocol. Serum and SR9011 concentrations are as indicated. The EdU+% at the 30-hr (D) and the E2f-ON% at the 24-hr (E) after serum stimulation are indicated to the right of individual histograms. Arrows indicate the approximate serum concentrations leading to EdU+% (D) and E2f-ON% (E) = 45%, respectively.

**Figure S4.**
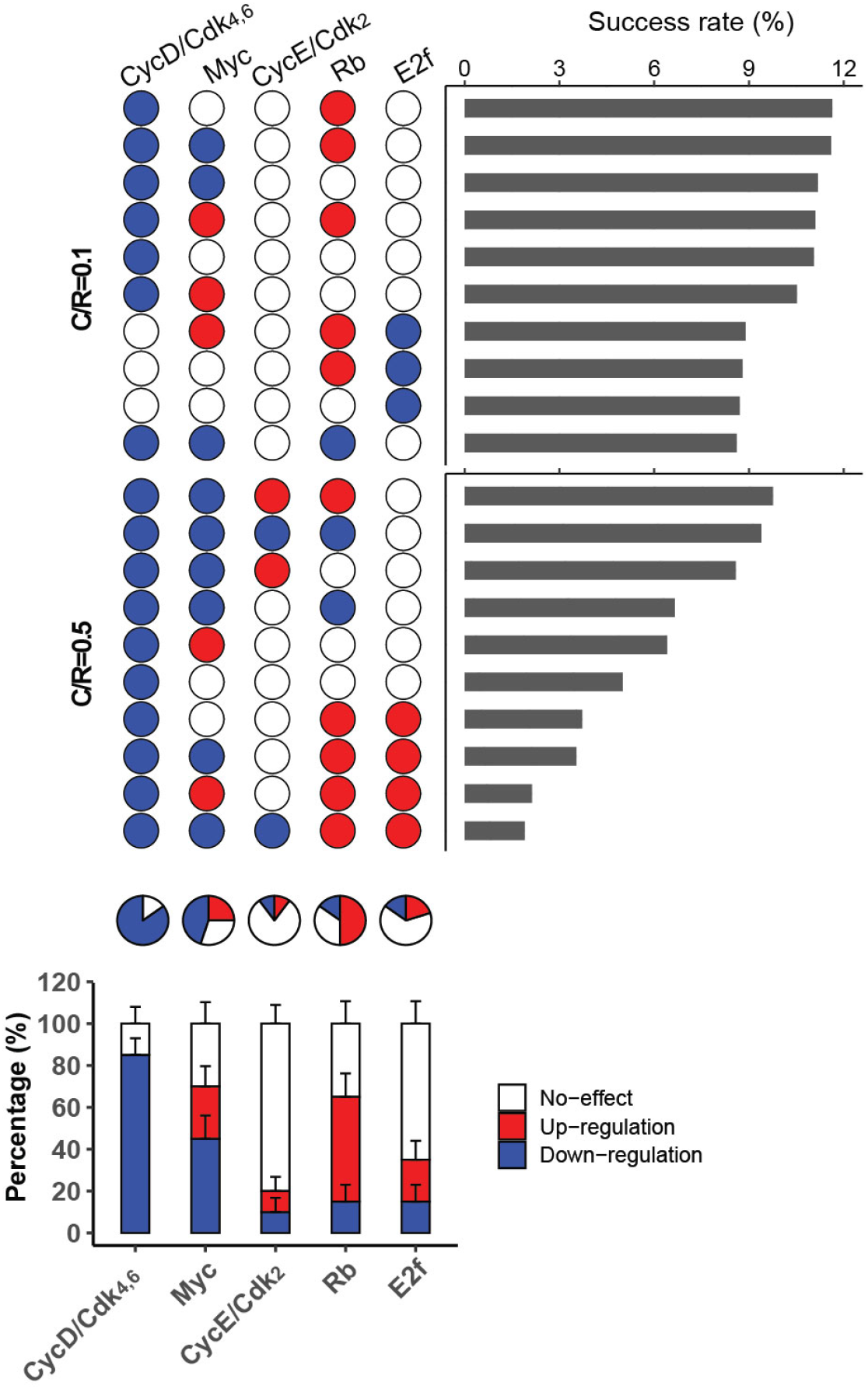
Link features of the 10 top-ranked models at each C/R level. (Top) The top 10 models at C/R = 0.1 and 0.5, respectively. The link features in each model are shown according to the upregulation (red), downregulation (blue), or no-effect (white) of the indicated target node by C/R. (Middle) Pie chart of the percentage of each link feature at the indicated target node among the combined 20 models (top). (Bottom) The average percentage of each link feature at the indicated target node between the two model groups (C/R = 0.1 and 0.5, respectively, top). Error bar, SD.

**Figure S5.**
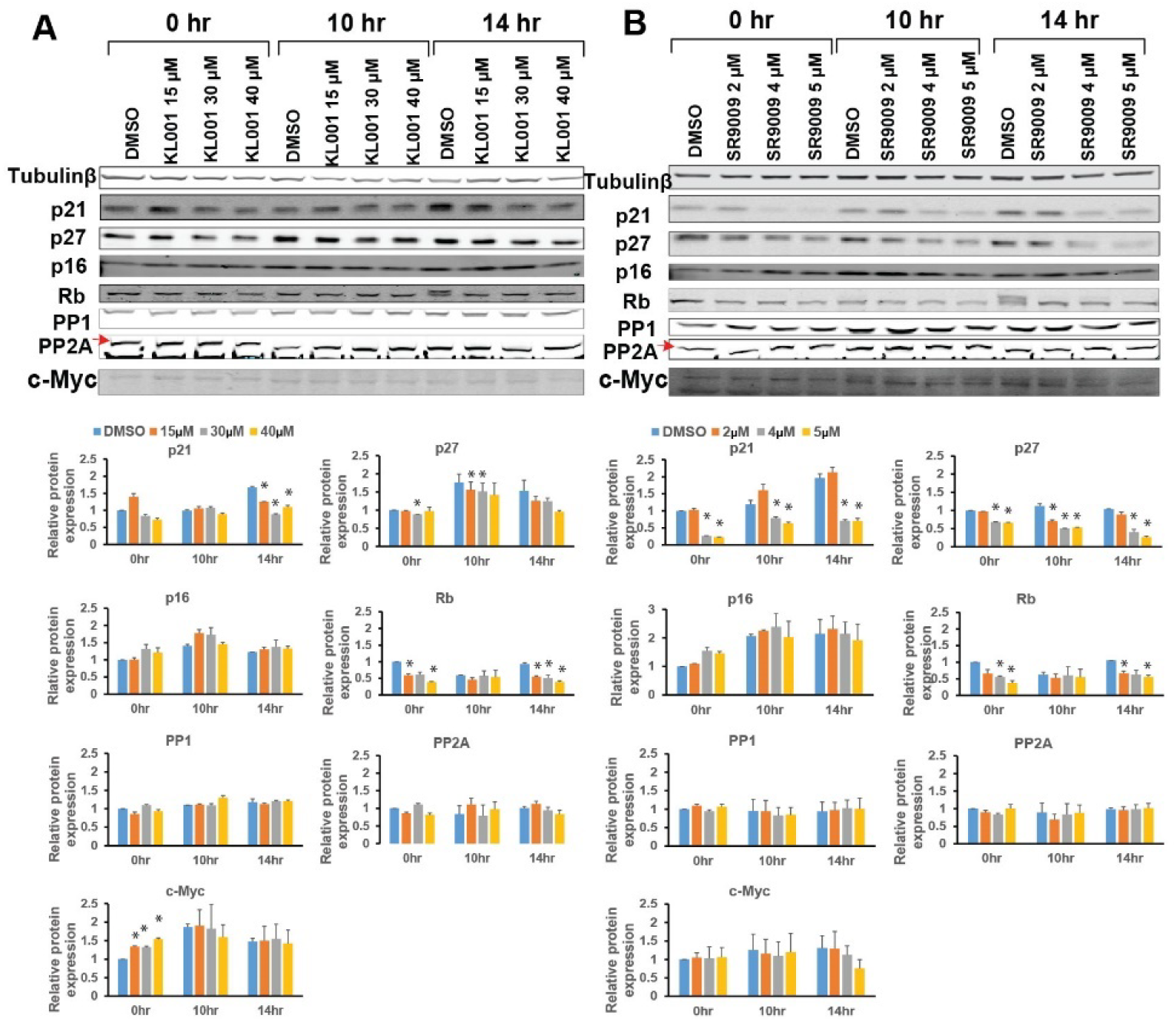
Western blot analysis of multiple Rb-E2f network proteins in response to Cry and Rev-erb agonists. The responses of indicated Rb-E2f network proteins to Cry agonist KL001 (A) and Rev-erb agonist SR9009 (B) at the indicated concentrations were measured following the STI.3dq/agonist protocol. Protein levels were measured by Western blot in quiescence (0 hr) and during cell cycle entry (10 and 14 hr after stimulated with 4% serum) and normalized to the 0-hr DMSO control. Error bar, SEM (n=2); *p < 0.05.

**Figure S6.**
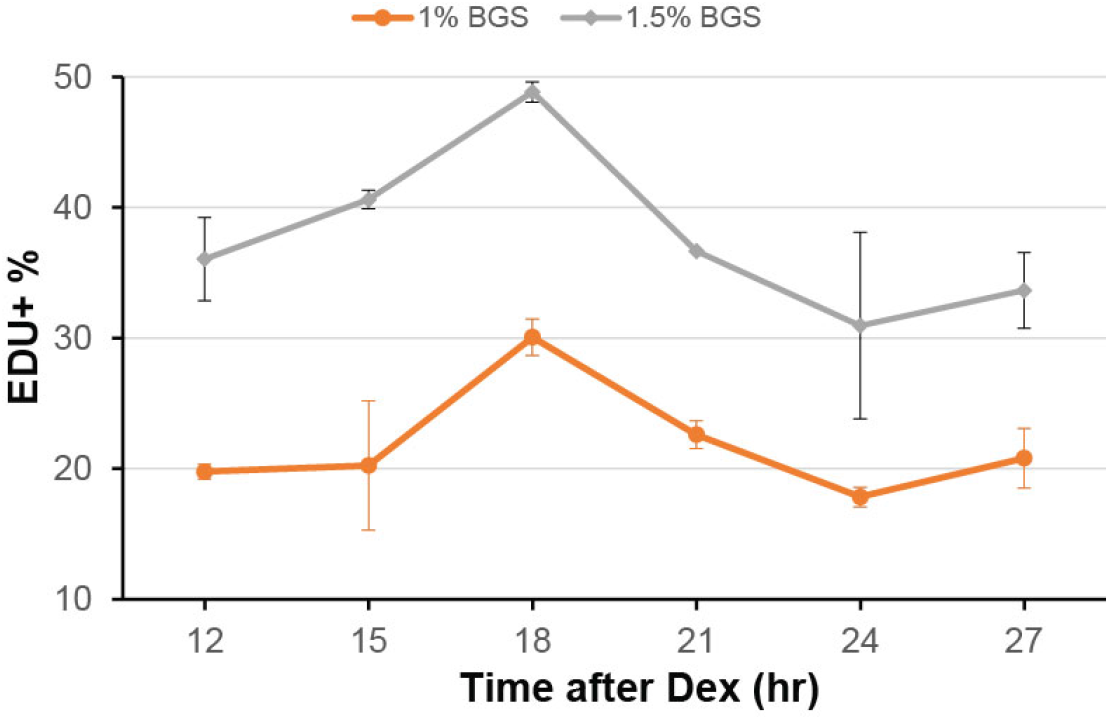
Circadian changes of quiescence depth. Quiescent (2-day serum-starved) cells were circadian-synchronized with dexamethasone (Dex, 100 nM) treatment for 2 hours followed by 12 hours of stabilization time (as in ^67^). Cells were subsequently, and every 3 hours afterward, stimulated with serum (1% and 1.5%, respectively) for 30 hours, followed by the measurement of EdU+%. Error bar, SEM (n⩾2).

**Table S1.** Rb-E2f bistable switch model with circadian regulation.

**Table S2.** Model parameters.

**Table S3.** Ranking for the top 10 topologies at the low level of C/R (= 0.1).

**Table S4.** Ranking for the top 10 topologies at the high level of C/R (= 0.5).

**Table S5.** Estimation of CycD/Cdk4,6 activity under KL001 and SR9009 treatments.

**Table S1.**
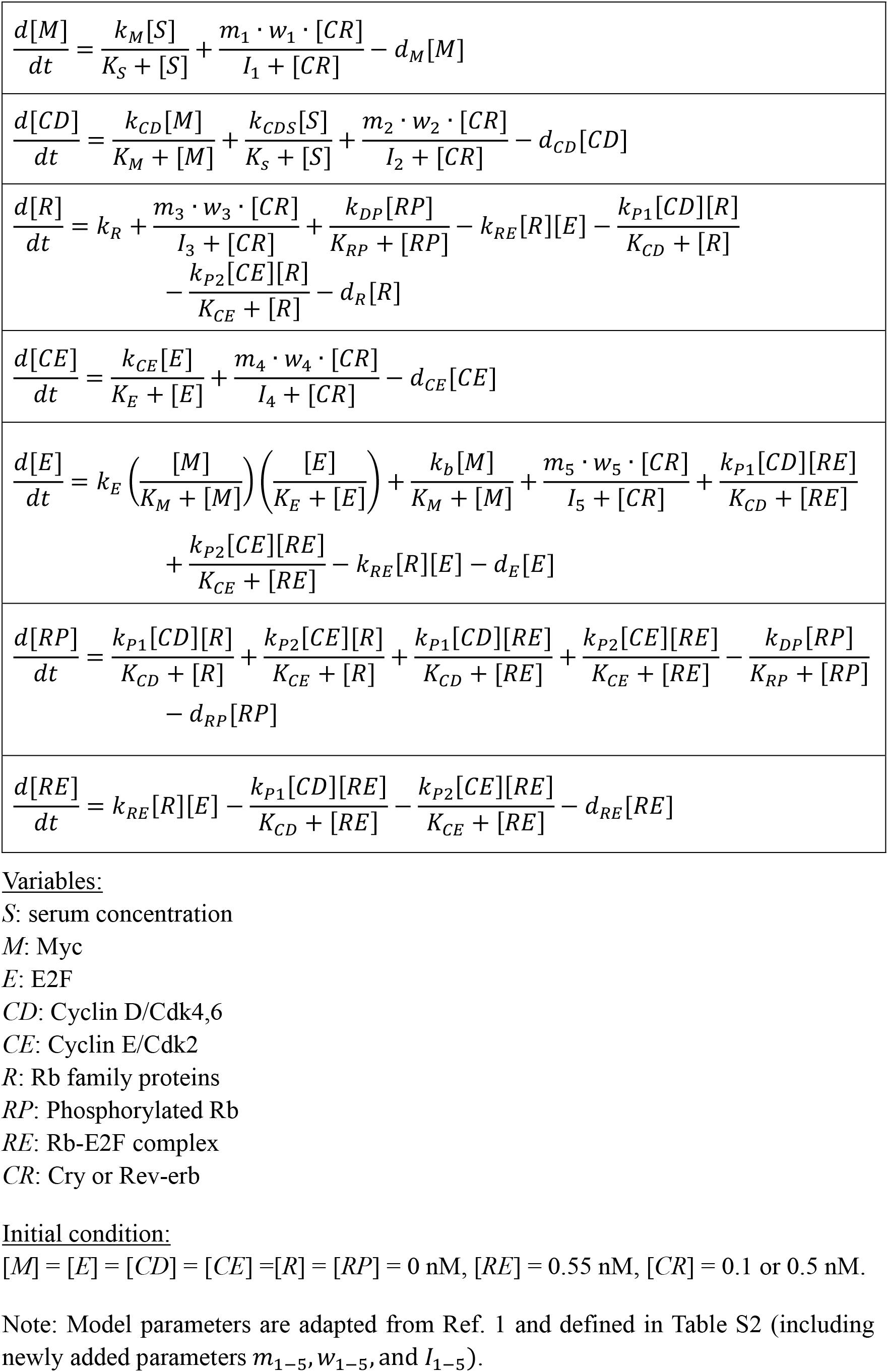
The Rb-E2f bistable switch model with circadian regulation (adapted from Ref. 1).

**Table S2.** Model parameters (adapted from Ref. 1)

| Symbol | Values | Description |
| --- | --- | --- |
| $k_M$ | 1.0 nM hr<sup>-1</sup> | Rate constant of Myc synthesis driven by growth factors |
| $k_E$ | 0.4 nM hr<sup>-1</sup> | Rate constant of E2F synthesis driven by Myc and E2F |
| $k_b$ | 0.003 nM hr<sup>-1</sup> | Rate constant of E2F synthesis driven by Myc alone |
| $k_{CD}$ | 0.03 nM hr<sup>-1</sup> | Rate constant of CycD synthesis driven by Myc |
| $k_{CDS}$ | 0.45 nM hr<sup>-1</sup> | Rate constant of CycD synthesis driven by growth factors |
| $k_{CE}$ | 0.35 nM hr<sup>-1</sup> | Rate constant of CycE synthesis driven by E2F |
| $k_R$ | 0.18 nM hr<sup>-1</sup> | Rate constant of Rb constitutive synthesis |
| $k_{P1}$ | 18 hr<sup>-1</sup> | Phosphorylation rate constant of Rb by CycD/Cdk4,6 |
| $k_{P2}$ | 18 hr<sup>-1</sup> | Phosphorylation rate constant of Rb by CycE/Cdk2 |
| $k_{DP}$ | 3.6 nM hr<sup>-1</sup> | Dephosphorylation rate constant of Rb by phosphatases |
| $k_{RE}$ | 180 nM<sup>-1</sup> hr<sup>-1</sup> | Association rate constant of Rb and E2F |
| $K_S$ | 0.5 nM | Michaelis-Menten parameter for CycD and Myc synthesis by growth factors |
| $K_E$ | 0.15 nM | Michaelis-Menten parameter for CycE and E2F synthesis by E2F |
| $K_M$ | 0.15 nM | Michaelis-Menten parameter for CycD and E2F synthesis by Myc |
| $K_{RP}$ | 0.01 nM | Michaelis-Menten parameter for Rb dephosphorylation |
| $K_{CD}$ | 0.92 nM | Michaelis-Menten parameter for Rb phosphorylation by CycD/Cdk4,6 |
| $K_{CE}$ | 0.92 nM | Michaelis-Menten parameter for Rb phosphorylation by CycE/Cdk2 |
| $d_M$ | 0.7 hr<sup>-1</sup> | Degradation rate constant of Myc |
| $d_E$ | 0.25 hr<sup>-1</sup> | Degradation rate constant of E2F |
| $d_{CD}$ | 1.5 hr<sup>-1</sup> | Degradation rate constant of CycD |
| $d_{CE}$ | 1.5 hr<sup>-1</sup> | Degradation rate constant of CycE |
| $d_R$ | 0.06 hr<sup>-1</sup> | Degradation rate constant of Rb |
| $d_{RP}$ | 0.06 hr<sup>-1</sup> | Degradation rate constant of phosphorylated Rb |
| $d_{RE}$ | 0.03 hr<sup>-1</sup> | Degradation rate constant of Rb-E2F complex |
| $m_1$ | 0, -1, or +1 | CR activate, inhibit, or have no effect on $M$ |
| $m_2$ | 0, -1, or +1 | CR activate, inhibit, or have no effect on $CD$ |
| $m_3$ | 0, -1, or +1 | CR activate, inhibit, or have no effect on $R$ |
| $m_4$ | 0, -1, or +1 | CR activate, inhibit, or have no effect on $CE$ |
| $m_5$ | 0, -1, or +1 | CR activate, inhibit, or have no effect on $E$ |
| $w_1$ | 1.0 nM hr<sup>-1</sup> | Rate constant of $CR$ effect on $M$ (matching the $k_m$ value) |
| $w_2$ | 0.45 nM hr<sup>-1</sup> | Rate constant of $CR$ effect on $CD$ (matching the $k_{CDS}$ value) |
| $w_3$ | 0.18 nM hr<sup>-1</sup> | Rate constant of $CR$ effect on $R$ (matching the $k_R$ value) |
| $w_4$ | 0.35 nM hr<sup>-1</sup> | Rate constant of $CR$ effect on $CE$ (matching the $k_{CE}$ value) |
| $w_5$ | 0.4 nM hr<sup>-1</sup> | Rate constant of $CR$ effect on $E$ (matching the $k_E$ value) |
| $I_1$ | 0.01 ~ 1 nM | Random Michaelis-Menten parameter for $CR$ effect on $M$ |
| $I_2$ | 0.01 ~ 1 nM | Random Michaelis-Menten parameter for $CR$ effect on $CD$ |
| $I_3$ | 0.01 ~ 1 nM | Random Michaelis-Menten parameter for $CR$ effect on $R$ |
| $I_4$ | 0.01 ~ 1 nM | Random Michaelis-Menten parameter for $CR$ effect on $CE$ |
| $I_5$ | 0.01 ~ 1 nM | Random Michaelis-Menten parameter for $CR$ effect on $E$ |

**Table S3.** Ranking of the top 10 topologies at the low level of C/R (= 0.1).

| Topology | 1250<br>Sim | 2500<br>Sim | 3750<br>Sim | 5000<br>Sim | 6250<br>Sim | 7500<br>Sim | 8750<br>Sim | 10,000<br>Sim |
| --- | --- | --- | --- | --- | --- | --- | --- | --- |
| #64 | 1 | 1 | 2 | 2 | 1 | 1 | 2 | 1 |
| #226 | 2 | 2 | 1 | 1 | 2 | 2 | 1 | 2 |
| #217 | 4 | 4 | 3 | 3 | 3 | 5 | 3 | 3 |
| #145 | 3 | 5 | 5 | 5 | 5 | 3 | 4 | 4 |
| #55 | 5 | 3 | 4 | 4 | 4 | 4 | 5 | 5 |
| #136 | 6 | 6 | 6 | 6 | 6 | 6 | 6 | 6 |
| #93 | 7 | 7 | 8 | 9 | 7 | 7 | 7 | 7 |
| #12 | 8 | 9 | 9 | 11 | 9 | 8 | 8 | 8 |
| #3 | 10 | 8 | 7 | 7 | 8 | 9 | 9 | 9 |
| #235 | 9 | 13 | 11 | 8 | 10 | 10 | 10 | 10 |

A total of 10,000 model simulations (Sim) were evenly divided into 8 bins (1250 Sim in each bin). The ranking of each topology (model number shown in the 1<sup>st</sup> column) was updated after each bin addition (i.e., after 1250, 2500, ..., 10,000 Sim).

**Table S4.** Ranking of the top 10 topologies at the high level of C/R (= 0.5).

| Topology | 1250<br>Sim | 2500<br>Sim | 3750<br>Sim | 5000<br>Sim | 6250<br>Sim | 7500<br>Sim | 8750<br>Sim | 10,000<br>Sim |
| --- | --- | --- | --- | --- | --- | --- | --- | --- |
| #229 | 2 | 2 | 1 | 1 | 1 | 1 | 1 | 1 |
| #241 | 1 | 1 | 2 | 2 | 2 | 2 | 2 | 2 |
| #220 | 3 | 3 | 3 | 3 | 3 | 3 | 3 | 3 |
| #235 | 5 | 5 | 5 | 5 | 5 | 4 | 4 | 4 |
| #136 | 4 | 4 | 4 | 4 | 4 | 5 | 5 | 5 |
| #55 | 6 | 6 | 6 | 6 | 6 | 6 | 6 | 6 |
| #65 | 8 | 7 | 7 | 7 | 7 | 7 | 7 | 7 |
| #227 | 7 | 8 | 8 | 8 | 8 | 8 | 8 | 8 |
| #146 | 10 | 9 | 10 | 9 | 9 | 9 | 9 | 9 |
| #233 | 9 | 10 | 11 | 10 | 10 | 10 | 10 | 10 |

See Table S3 legend for detail.

**Table S5.** Estimation of CycD/Cdk4,6 activity under KL001 and SR9009 treatments.

|  |  |  |  |  |  |  |  |  |  |  |  |  |  |
| --- | --- | --- | --- | --- | --- | --- | --- | --- | --- | --- | --- | --- | --- |
| **KL001** |  |  |  |  |  |  |  |  |  |  |  |  |  |
| treatment (rep 1) | CycD1 | CycD3 | Cdk4 | Cdk6 | p21 | p27 | p16 | JA (w/o adj. exp) | JA' (w/ adj. exp) | Mean JA' | sd | normalized JA' | sd |
| DMSO | 3.59 | 0.90 | 1.22 | 2.29 | 1.33 | 1.91 | 1.33 | 2.58 | 4.66 | 4.96 | 0.43 | **1.00** | 0.00 |
| KL001 15uM | 3.15 | 0.71 | 1.14 | 1.43 | 1.13 | 1.58 | 1.53 | 1.76 | 3.99 | 3.88 | 0.16 | **0.78** | 0.07 |
| KL001 30uM | 2.53 | 0.53 | 1.18 | 1.14 | 1.01 | 1.54 | 1.56 | 1.30 | 3.38 | 3.51 | 0.18 | **0.71** | 0.07 |
| KL001 40uM | 1.67 | 0.47 | 0.90 | 0.47 | 0.99 | 1.37 | 1.45 | 0.57 | 1.92 | 1.60 | 0.45 | **0.32** | 0.10 |
| treatment (rep 2) | CycD1 | CycD3 | CDK4 | CDK6 | p21 | p27 | p16 | JA (w/o adj. exp) | JA' (w/ adj. exp) |  |  |  |  |
| DMSO | 2.61 | 1.23 | 1.42 | 2.29 | 1.34 | 1.39 | 1.29 | 2.66 | 5.26 |  |  |  |  |
| KL001 15uM | 1.94 | 0.95 | 1.32 | 1.72 | 1.18 | 1.23 | 1.55 | 1.66 | 3.77 |  |  |  |  |
| KL001 30uM | 1.30 | 0.94 | 1.42 | 1.25 | 0.94 | 1.21 | 1.55 | 1.21 | 3.63 |  |  |  |  |
| KL001 40uM | 0.78 | 0.75 | 0.67 | 0.48 | 0.99 | 1.00 | 1.33 | 0.40 | 1.28 |  |  |  |  |
| **SR9009** |  |  |  |  |  |  |  |  |  |  |  |  |  |
| treatment (rep 1) | CycD1 | CycD3 | Cdk4 | Cdk6 | p21 | p27 | p16 | JA (w/o adj. exp) | JA' (w/ adj. exp) | Mean JA' | sd | normalized JA' | sd |
| DMSO | 4.10 | 1.27 | 1.53 | 1.59 | 1.45 | 1.11 | 2.39 | 2.53 | 6.03 | 5.77 | 0.37 | **1.00** | 0.00 |
| SR9009 2uM | 4.65 | 0.97 | 1.12 | 1.15 | 1.71 | 0.85 | 2.52 | 1.89 | 4.17 | 3.95 | 0.30 | **0.69** | 0.07 |
| SR9009 4uM | 0.47 | 0.37 | 0.56 | 0.44 | 0.70 | 0.42 | 2.70 | 0.16 | 0.52 | 0.50 | 0.02 | **0.09** | 0.01 |
| SR9009 5uM | 0.22 | 0.31 | 0.52 | 0.32 | 0.61 | 0.38 | 2.54 | 0.09 | 0.34 | 0.31 | 0.05 | **0.06** | 0.01 |
| treatment (rep 2) | CycD1 | CycD3 | CDK4 | CDK6 | p21 | p27 | p16 | JA (w/o adj. exp) | JA' (w/ adj. exp) |  |  |  |  |
| DMSO | 3.83 | 1.20 | 1.46 | 1.36 | 1.69 | 1.06 | 1.82 | 2.33 | 5.50 |  |  |  |  |
| SR9009 2uM | 4.45 | 0.91 | 1.08 | 0.92 | 2.03 | 0.74 | 2.03 | 1.68 | 3.74 |  |  |  |  |
| SR9009 4uM | 0.49 | 0.28 | 0.52 | 0.26 | 0.78 | 0.49 | 1.84 | 0.14 | 0.49 |  |  |  |  |
| SR9009 5uM | 0.20 | 0.16 | 0.49 | 0.21 | 0.72 | 0.42 | 1.41 | 0.07 | 0.27 |  |  |  |  |
| *Gene expression (estimated from Ref [2])* |  |  |  |  |  |  |  |  |  |  |  |  |  |
|  | CycD1 | CycD3 | Cdk4 | Cdk6 | p21 | p27 | p16 |  |  |  |  |  |  |
| relative expression | 0.64 | 1.00 | 0.89 | 0.03 | 0.30 | 0.10 | 0.15 |  |  |  |  |  |  |
| sd | 0.00013 | 0 | 0.00023 | 2.2E-05 | 0.00012 | 2.4E-05 | 8.1E-05 |  |  |  |  |  |  |

The protein levels of CycD1, CycD3, Cdk4, Cdk6, p21, p27, and p16 were derived from Western blot as in Fig. 3 and S5 and averaged over 10 and 14 hr in each of the two replicates (rep 1 and 2). The joint activity (JA) of CycD/Cdk4,6 was assumed to be positively proportional to the combined levels of CycD and Cdk4,6 components and negatively proportional to the combined level of CKI components, JA = (CycD1+CycD3)(Cdk4+Cdk6)/(p16+p21+p27). Next, JA was adjusted (JA’) according to the relative gene expression level of each component (estimated from Ref. 2 based on the RNA-seq data in growing REF/E23 cells). The mean JA’ value was obtained by averaging over rep 1 and 2 at each treatment condition and subsequently normalized to the DMSO control. (sd, standard deviation).

## Notes

### Competing Interest Statement

The authors have declared no competing interest.

## REFERENCES

1. Coller, H.A., Sang, L. & Roberts, J.M. A new description of cellular quiescence. PLoS Biol 4, e83 (2006).

2. Cheung, T.H. & Rando, T.A. Molecular regulation of stem cell quiescence. Nat Rev Mol Cell Biol 14, 329–340 (2013).

3. Rumman, M., Dhawan, J. & Kassem, M. Concise Review: Quiescence in Adult Stem Cells: Biological Significance and Relevance to Tissue Regeneration. Stem Cells 33, 2903–2912 (2015).

4. Bouchard-Cannon, P., Mendoza-Viveros, L., Yuen, A., Kaern, M. & Cheng, H.Y. The circadian molecular clock regulates adult hippocampal neurogenesis by controlling the timing of cell-cycle entry and exit. Cell Rep 5, 961–973 (2013).

5. Gengatharan, A. et al. Adult neural stem cell activation in mice is regulated by the day/night cycle and intracellular calcium dynamics. Cell 184, 709–722. e713 (2021).

6. Janich, P. et al. The circadian molecular clock creates epidermal stem cell heterogeneity. Nature 480, 209–214 (2011).

7. Panda, S. et al. Coordinated transcription of key pathways in the mouse by the circadian clock. Cell 109, 307–320 (2002).

8. Ueda, H.R. et al. A transcription factor response element for gene expression during circadian night. Nature 418, 534–539 (2002).

9. Storch, K.F. et al. Extensive and divergent circadian gene expression in liver and heart. Nature 417, 78–83 (2002).

10. Sancar, A. & Van Gelder, R.N. Clocks, cancer, and chronochemotherapy. Science 371 (2021).

11. Takahashi, J.S. Transcriptional architecture of the mammalian circadian clock. Nat Rev Genet 18, 164–179 (2017).

12. Hong, C.I. et al. Circadian rhythms synchronize mitosis in Neurospora crassa. Proc Natl Acad Sci U S A 111, 1397–1402 (2014).

13. Yang, Q., Pando, B.F., Dong, G., Golden, S.S. & van Oudenaarden, A. Circadian gating of the cell cycle revealed in single cyanobacterial cells. Science 327, 1522–1526 (2010).

14. Bieler, J. et al. Robust synchronization of coupled circadian and cell cycle oscillators in single mammalian cells. Mol Syst Biol 10, 739 (2014).

15. Feillet, C. et al. Phase locking and multiple oscillating attractors for the coupled mammalian clock and cell cycle. Proc Natl Acad Sci U S A 111, 9828–9833 (2014).

16. Liu, Z. et al. Circadian regulation of c-MYC in mice. Proc Natl Acad Sci U S A 117, 21609–21617 (2020).

17. Fu, L., Pelicano, H., Liu, J., Huang, P. & Lee, C. The circadian gene Period2 plays an important role in tumor suppression and DNA damage response in vivo. Cell 111, 41–50 (2002).

18. Gréchez-Cassiau, A., Rayet, B., Guillaumond, F., Teboul, M. & Delaunay, F. The circadian clock component BMAL1 is a critical regulator of p21WAF1/CIP1 expression and hepatocyte proliferation. Journal of Biological Chemistry 283, 4535–4542 (2008).

19. Matsuo, T. et al. Control mechanism of the circadian clock for timing of cell division in vivo. Science 302, 255–259 (2003).

20. Sahar, S. & Sassone-Corsi, P. Metabolism and cancer: the circadian clock connection. Nat Rev Cancer 9, 886–896 (2009).

21. Kwon, J.S. et al. Controlling Depth of Cellular Quiescence by an Rb-E2F Network Switch. Cell Rep 20, 3223–3235 (2017).

22. Brooks, R.F., Richmond, F.N., Riddle, P.N. & Richmond, K.M. Apparent heterogeneity in the response of quiescent swiss 3T3 cells to serum growth factors: implications for the transition probability model and parallels with “cellular senescence” and “competence”. J Cell Physiol 121, 341–350 (1984).

23. Fujimaki, K. et al. Graded regulation of cellular quiescence depth between proliferation and senescence by a lysosomal dimmer switch. Proceedings of the National Academy of Sciences 116, 22624–22634 (2019).

24. Augenlicht, L.H. & Baserga, R. Changes in the G0 state of WI-38 fibroblasts at different times after confluence. Exp Cell Res 89, 255–262 (1974).

25. Bucher, N.L. Regeneration of Mammalian Liver. Int Rev Cytol 15, 245–300 (1963).

26. Roth, G.S. & Adelman, R.C. Age-dependent regulation of mammalian DNA synthesis and cell division in vivo by glucocorticoids. Exp Gerontol 9, 27–31 (1974).

27. Yanez, I. & O’Farrell, M. Variation in the length of the lag phase following serum restimulation of mouse 3T3 cells. Cell Biol Int Rep 13, 453–462 (1989).

28. Rodgers, J.T. et al. mTORC1 controls the adaptive transition of quiescent stem cells from G0 to G(Alert). Nature 510, 393–396 (2014).

29. Llorens-Bobadilla, E. et al. Single-Cell Transcriptomics Reveals a Population of Dormant Neural Stem Cells that Become Activated upon Brain Injury. Cell Stem Cell 17, 329–340 (2015).

30. Rodgers, J.T., Schroeder, M.D., Ma, C. & Rando, T.A. HGFA is an injury-regulated systemic factor that induces the transition of stem cells into GAlert. Cell reports 19, 479–486 (2017).

31. Lee, G. et al. Fully reduced HMGB1 accelerates the regeneration of multiple tissues by transitioning stem cells to GAlert. Proc Natl Acad Sci U S A 115, E4463–E4472 (2018).

32. Yao, G., Lee, T.J., Mori, S., Nevins, J.R. & You, L. A bistable Rb-E2F switch underlies the restriction point. Nat Cell Biol 10, 476–482 (2008).

33. Wang, X. et al. Exit from quiescence displays a memory of cell growth and division. Nat Commun 8, 321 (2017).

34. Wu, L. et al. The E2F1-3 transcription factors are essential for cellular proliferation. Nature 414, 457–462 (2001).

35. Johnson, D.G., Schwarz, J.K., Cress, W.D. & Nevins, J.R. Expression of transcription factor E2F1 induces quiescent cells to enter S phase. Nature 365, 349–352 (1993).

36. Yao, G. Modelling mammalian cellular quiescence. Interface Focus 4, 20130074 (2014).

37. Cappell, S.D., Chung, M., Jaimovich, A., Spencer, S.L. & Meyer, T. Irreversible APCCdh1 inactivation underlies the point of no return for cell-cycle entry. Cell 166, 167–180 (2016).

38. Cappell, S.D. et al. EMI1 switches from being a substrate to an inhibitor of APC/C CDH1 to start the cell cycle. Nature 558, 313–317 (2018).

39. Leung, J.Y., Ehmann, G.L., Giangrande, P.H. & Nevins, J.R. A role for Myc in facilitating transcription activation by E2F1. Oncogene 27, 4172–4179 (2008).

40. Hirota, T. et al. Identification of small molecule activators of cryptochrome. Science 337, 1094–1097 (2012).

41. Calzone, L., Gelay, A., Zinovyev, A., Radvanyi, F. & Barillot, E. A comprehensive modular map of molecular interactions in RB/E2F pathway. Mol Syst Biol 4, 173 (2008).

42. Sears, R.C. & Nevins, J.R. Signaling networks that link cell proliferation and cell fate. Journal of Biological Chemistry 277, 11617–11620 (2002).

43. Attwooll, C., Lazzerini Denchi, E. & Helin, K. The E2F family: specific functions and overlapping interests. EMBO J 23, 4709–4716 (2004).

44. Frolov, M.V. & Dyson, N.J. Molecular mechanisms of E2F-dependent activation and pRB-mediated repression. J Cell Sci 117, 2173–2181 (2004).

45. Weinberg, R.A. The retinoblastoma protein and cell cycle control. Cell 81, 323–330 (1995).

46. Yao, G., Tan, C., West, M., Nevins, J.R. & You, L. Origin of bistability underlying mammalian cell cycle entry. Mol Syst Biol 7, 485 (2011).

47. Ma, W., Lai, L., Ouyang, Q. & Tang, C. Robustness and modular design of the Drosophila segment polarity network. Mol Syst Biol 2, 70 (2006).

48. Ma, W., Trusina, A., El-Samad, H., Lim, W.A. & Tang, C. Defining network topologies that can achieve biochemical adaptation. Cell 138, 760–773 (2009).

49. Arora, M., Moser, J., Phadke, H., Basha, A.A. & Spencer, S.L. Endogenous Replication Stress in Mother Cells Leads to Quiescence of Daughter Cells. Cell Rep 19, 1351–1364 (2017).

50. Barr, A.R. et al. DNA damage during S-phase mediates the proliferation-quiescence decision in the subsequent G1 via p21 expression. Nat Commun 8, 14728 (2017).

51. Laurenti, E. et al. CDK6 levels regulate quiescence exit in human hematopoietic stem cells. Cell Stem Cell 16, 302–313 (2015).

52. Yang, H.W., Chung, M., Kudo, T. & Meyer, T. Competing memories of mitogen and p53 signalling control cell-cycle entry. Nature 549, 404–408 (2017).

53. Min, M., Rong, Y., Tian, C. & Spencer, S.L. Temporal integration of mitogen history in mother cells controls proliferation of daughter cells. Science 368, 1261–1265 (2020).

54. Narasimha, A.M. et al. Cyclin D activates the Rb tumor suppressor by mono-phosphorylation. Elife 3 (2014).

55. Pennycook, B.R. & Barr, A.R. Restriction point regulation at the crossroads between quiescence and cell proliferation. FEBS letters 594, 2046–2060 (2020).

56. Schwarz, C. et al. A precise Cdk activity threshold determines passage through the restriction point. Molecular cell 69, 253–264. e255 (2018).

57. Stallaert, W., Kedziora, K.M., Chao, H.X. & Purvis, J.E. Bistable switches as integrators and actuators during cell cycle progression. FEBS letters 593, 2805–2816 (2019).

58. Brooks, R.F. Cell cycle commitment and the origins of cell cycle variability. Frontiers in Cell and Developmental Biology 9, 1891 (2021).

59. Matson, J.P. & Cook, J.G. Cell cycle proliferation decisions: the impact of single cell analyses. FEBS J 284, 362–375 (2017).

60. Novák, B. & Tyson, J.J. Mechanisms of signalling-memory governing progression through the eukaryotic cell cycle. Current Opinion in Cell Biology 69, 7–16 (2021).

61. Goldbeter, A. & Koshland, D.E., Jr. An amplified sensitivity arising from covalent modification in biological systems. Proc Natl Acad Sci U S A 78, 6840–6844 (1981).

62. Overton, K.W., Spencer, S.L., Noderer, W.L., Meyer, T. & Wang, C.L. Basal p21 controls population heterogeneity in cycling and quiescent cell cycle states. Proc Natl Acad Sci U S A 111, E4386–4393 (2014).

63. Hoops, S. et al. COPASI: a COmplex PAthway SImulator. Bioinformatics 22, 3067–3074 (2006).

64. Lee, T.J., Yao, G., Bennett, D.C., Nevins, J.R. & You, L. Stochastic E2F activation and reconciliation of phenomenological cell-cycle models. PLoS Biol 8 (2010).

65. Gillespie, D.T. The chemical Langevin equation. The Journal of Chemical Physics 113, 297–306 (2000).

66. Dengler, W.A., Schulte, J., Berger, D.P., Mertelsmann, R. & Fiebig, H.H. Development of a propidium iodide fluorescence assay for proliferation and cytotoxicity assays. Anticancer Drugs 6, 522–532 (1995).

67. Phillips, N.E. et al. The circadian oscillator analysed at the single-transcript level. Molecular systems biology 17, e10135 (2021).

## Supplementary References

1. Yao, G., Lee, T.J., Mori, S., Nevins, J.R. & You, L. A bistable Rb-E2F switch underlies the restriction point. Nat Cell Biol 10, 476–482 (2008).

2. Fujimaki, K. et al. Graded regulation of cellular quiescence depth between proliferation and senescence by a lysosomal dimmer switch. Proceedings of the National Academy of Sciences 116, 22624–22634 (2019).

